# Structural basis for the neutralization of SARS-CoV-2 by an antibody from a convalescent patient

**DOI:** 10.1101/2020.06.12.148387

**Authors:** Daming Zhou, Helen ME Duyvesteyn, Cheng-Pin Chen, Chung-Guei Huang, Ting-Hua Chen, Shin-Ru Shih, Yi-Chun Lin, Chien-Yu Cheng, Shu-Hsing Cheng, Yhu-Chering Huang, Tzou-Yien Lin, Che Ma, Jiandong Huo, Loic Carrique, Tomas Malinauskas, Reinis R Ruza, Pranav NM Shah, Tiong Kit Tan, Pramila Rijal, Robert F. Donat, Kerry Godwin, Karen Buttigieg, Julia Tree, Julika Radecke, Neil G Paterson, Piyasa Supasa, Juthathip Mongkolsapaya, Gavin R Screaton, Miles W. Carroll, Javier G. Jaramillo, Michael Knight, William James, Raymond J Owens, James H. Naismith, Alain Townsend, Elizabeth E Fry, Yuguang Zhao, Jingshan Ren, David I Stuart, Kuan-Ying A. Huang

**Affiliations:** Division of Structural Biology, University of Oxford, The Wellcome Centre for Human Genetics, Headington, Oxford, OX3 7BN, UK; Department of Infectious Diseases, Taoyuan General Hospital, Ministry of Health and Welfare, Taoyuan, and National Yang-Ming University, Taipei, Taiwan; Research Center for Emerging Viral Infections, College of Medicine, Chang Gung University, Taoyuan, Taiwan; Department of Laboratory Medicine, Chang Gung Memorial Hospital, Taoyuan, Taiwan; Genomics Research Center, Academia Sinica, Taipei, Taiwan; Department of Infectious Diseases, Taoyuan General Hospital, Ministry of Health and Welfare, Taoyuan, and Taipei Medical University, Taipei, Taiwan; Division of Pediatric Infectious Diseases, Department of Pediatrics, Chang Gung Memorial Hospital, Taoyuan, Taiwan; The Rosalind Franklin Institute, Harwell Campus, OX11 0FA, UK; Protein Production UK, Research Complex at Harwell, Harwell Science & Innovation Campus, Didcot, OX11 0FA, UK; MRC Human Immunology Unit, Weatherall Institute of Molecular Medicine, University of Oxford, John Radcliffe Hospital, Oxford, OX3 9DS, UK; Centre for Translational Immunology, Chinse Academy of Medical Sciences Oxford Institute, University of Oxford, Oxford OX3 7FZ, UK; National Infection Service, Public Health England, Porton Down, Salisbury, SP4 0JG, UK; Diamond Light Source Ltd, Harwell Science & Innovation Campus, Didcot, OX11 0DE, UK; Nuffield Department of Medicine, Wellcome Centre for Human Genetics, University of Oxford, Oxford, UK; Dengue Hemorrhagic Fever Research Unit, Office for Research and Development, Faculty of Medicine, Siriraj Hospital, Mahidol University, Bangkok, Thailand; William Dunn School of Pathology, University of Oxford, South Parks Road, Oxford, OX1 3RE, UK

## Abstract

The COVID-19 pandemic has had unprecedented health and economic impact, but currently there are no approved therapies. We have isolated an antibody, EY6A, from a late-stage COVID-19 patient and show it neutralises SARS-CoV-2 and cross-reacts with SARS-CoV-1. EY6A Fab binds tightly (K_D_ of 2 nM) the receptor binding domain (RBD) of the viral Spike glycoprotein and a 2.6Å crystal structure of an RBD/EY6A Fab complex identifies the highly conserved epitope, away from the ACE2 receptor binding site. Residues of this epitope are key to stabilising the pre-fusion Spike. Cryo-EM analyses of the pre-fusion Spike incubated with EY6A Fab reveal a complex of the intact trimer with three Fabs bound and two further multimeric forms comprising destabilized Spike attached to Fab. EY6A binds what is probably a major neutralising epitope, making it a candidate therapeutic for COVID-19.

---

SARS-CoV-2 emerged in December 2019 resulting in a pandemic with an estimated 5-6 % mortality rate^1^. Akin to SARS-CoV-1, the causative agent of the 2003 SARS outbreak, this is an enveloped beta-coronavirus with protrusions of large trimeric ‘Spike’ proteins. Receptor binding domains (RBD) located at the tip of these Spikes facilitate host cell entry via interaction with angiotensin-converting enzyme 2 (ACE2). Spikes are type I transmembrane glycoproteins, formed from a single polypeptide, which transitions into a post-fusion state via cleavage into S1 (N-terminal) and S2 (C-terminal) chains following receptor binding or trypsin treatment. In the pre-fusion state, the apical RBD (~22 kDa) is folded down enshrouded by the N-terminal domain of the Spike so that the receptor binding site is inaccessible until, it is assumed, an RBD stochastically swings upwards to present the ACE2 binding site^2–5^. ACE2 interaction locks the RBD in the ‘up’ conformation which drives conversion to the post-fusion form where the S2 subunit engages the host membrane whilst dispensing with S1^2,3^

Neutralising human monoclonal antibodies that recognise the ACE2 receptor binding site for SARS-CoV-1 and SARS-CoV-2 are generally not cross-reactive between the two viruses and are susceptible to escape mutation^6–10^ (indeed a natural mutation Y495N has already been identified at this site (GISAID^11^: Accession ID: EPI_ISL_429783 Wienecke-Baldacchino et al.)). In contrast CR3022 (derived from a SARS-CoV-1 patient) cross-reacts strongly with SARS-CoV-2 (Methods, Fig. 1) and has been shown to recognise a cryptic, conserved epitope on the RBD distinct from the binding epitope of ACE2^7,12–14^. That this is not uncommon for SARS-CoV-1 antibodies is suggested by similar observations for 47D11^15^.

**Fig. 1.**
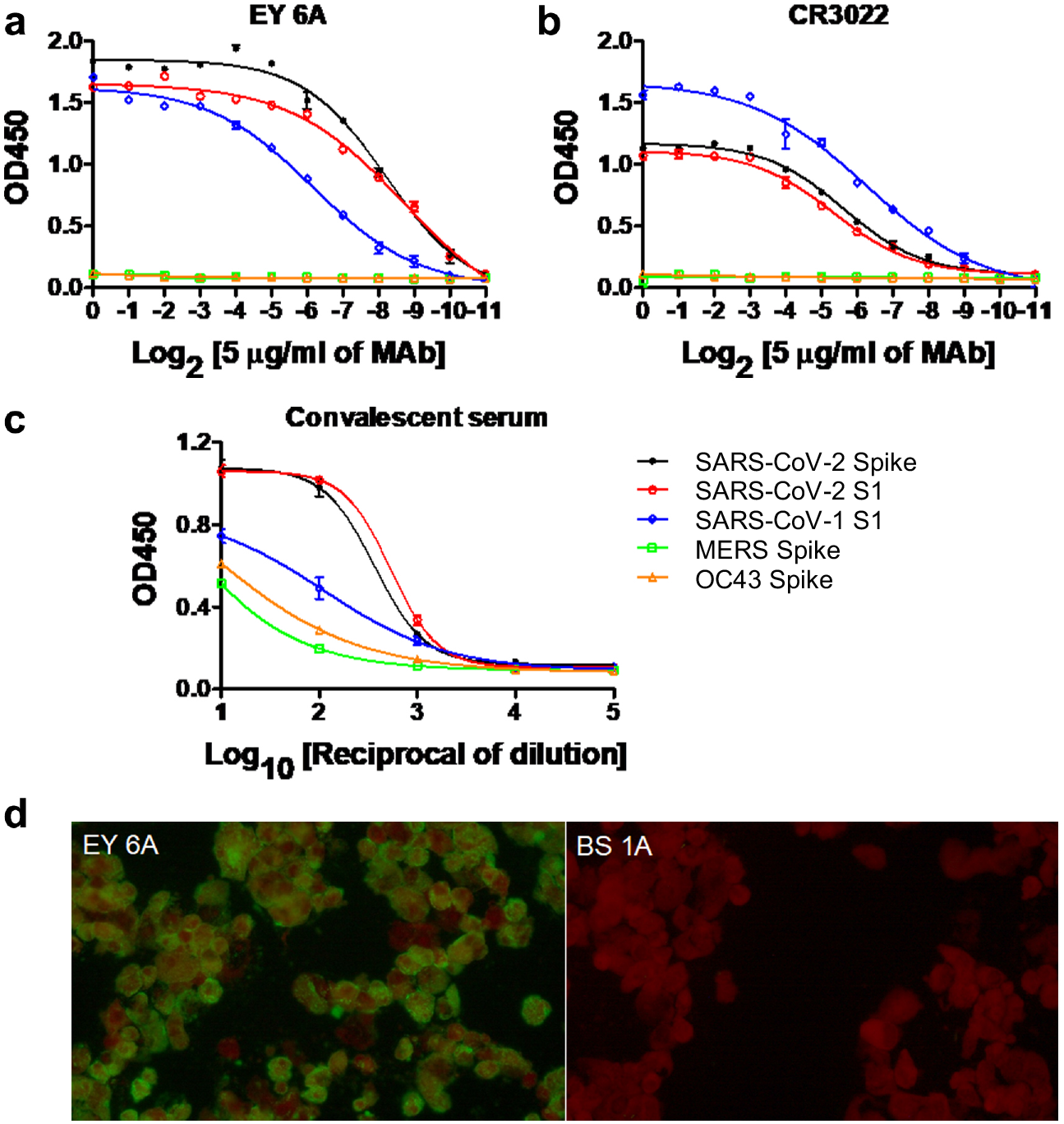
Binding specificity of EY6A in ELISA and immunofluorescence. **a,** Antibody EY6A binds the S1 subunit of SARS-CoV-2 and cross react with S1 of SARS-CoV-1. **b**, Antibody CR3022 similarly binds the S1 subunit of SARS-CoV-2 and cross react with SARS-CoV-1 S1, but with lower affinity. **c**, Convalescent serum from a COVID-19 patient was also included as a control and in this case binding to MERS and OC43 Spike proteins also investigated. **d**, Binding of EY6A on the SARS-CoV-2 infected cells in immunofluorescence assay. Anti-influenza H3 MAb BS 1A was included as a control.

## Results

### EY6A binds Spike RBD with little effect on ACE2 binding

To isolate SARS-CoV-2 Spike-reactive monoclonal antibodies, we cloned antibody genes from blood-derived plasmablasts of a COVID-19 patient in the convalescent phase. One of these, EY6A was shown by ELISA to bind S1 of SARS-CoV-2 and cross react with SARS-CoV-1 (Fig. 1). Binding of EY6A to SARS-CoV-2-infected cells was detected by immunofluorescence (Fig. 1). Surface plasmon resonance (SPR) measurements for EY6A Fab showed high affinity binding to immobilised SARS-CoV-2 RBD (K_D_ = 2 nM, although the value for immobilised EY6A IgG was somewhat higher) as derived from the kinetic data (Methods, Extended Data Fig. 1, Extended Data Table 1). SPR studies showed that there was some interdependence of EY6A and CR3022 binding, which varied depending on which component was immobilised on to the sensor chip; ACE2 blocking assays confirmed a somewhat asymmetric blocking effect (Extended Data Fig. 2). With RBD stably expressed on MDCK-SIAT1 cells (MDCK-RBD), EY6A did not block binding of ACE2 to the RBD, whereas with ACE2 stably expressed on MDCK-SIAT1 cells (MDCK-ACE2) EY6A blocked the interaction of RBD with ACE2. In this assay, EY6A exhibits around 7 times stronger ACE2 blocking than CR3022^13^ (EY6A, IC50 = 54 nM; CR3022, IC50 = 347 nM) and has equivalent ACE2 inhibition compared to ACE2-Fc (IC50 = 54 nM) and VHH72-Fc (IC50 = 69 nM)^2^. These observations are suggestive of an indirect effect by EY6A once bound to the RBD, consistent with an allosteric or weak direct interaction. This was supported by an SPR competition assay with immobilised CR3022, which binds distant from the ACE2 binding site (Extended Data Fig. 2)^12^. This showed complete competition with EY6A for RBD binding suggesting they recognise the same or overlapping epitopes, and indicated that EY6A binds the SARS-CoV-2 RBD more tightly.

### EY6A neutralises SARS-CoV-2

Two independent neutralisation tests, both using live wild type SARS-CoV-2 showed strong neutralisation. A neutralisation test for EY6A based on quantitative PCR detection of virus in the supernatant bathing infected Vero E6 cells after 5 days of culture, showed a ~1000-fold reduction in virus signal (Methods, Extended Data Fig. 3) indicating that it is highly neutralising. This was further corroborated by a plaque reduction neutralisation test (PRNT) at PHE Porton Down (Methods and Extended Data Table 2) using SARS-CoV-2 virus and EY6A which gave an ND_50_ of ~10.8 μg/mL (70 nM) (calculated according to Grist^16^). A separate PRNT implementation at Oxford gave a slightly higher ND_50_ of ~30 μg/mL, consistent with a shorter incubation time of antibody with virus at lower temperature (Extended Data Fig. 4).

### RBD/EY6A Fab complex reveals attachment to a conserved epitope

To elucidate the epitope of EY6A, we determined the crystal structures of the deglycosylated SARS-CoV-2 RBD in complex with EY6A Fab alone and in a ternary complex incorporating a nanobody (Nb) which has been shown to compete with ACE2 (for data on a closely related Nb see Huo 2020, submitted), as a crystallisation chaperone. The crystals of the binary complex diffracted to 3.8 Å resolution (Methods, Extended Data Table 3) and those of the ternary complex to 2.65 Å. The interaction between EY6A and the RBD was identical in both complexes (Extended Data Fig. 5). The higher resolution ternary complex, which showed that there was no interaction between EY6A and the Nb, permitted a full interpretation of the detailed interactions (Figs. 2 and 3) and has been refined to give an R-work/R-free of 0.216/0.262 and good stereochemistry (Methods, Extended Data Table 3). Residues 333-527 of the RBD, 1-136 and 141-224 of the heavy chain and 1-215 of the light chain of EY6A and 2-126 of the Nb are well defined Fig. 2a,b. The Nb recognises an epitope adjacent to and slightly overlapping the ACE2 receptor binding site and binds the RBD orthogonally to EY6A (Fig. 2b,c). EY6A binds essentially the same epitope as CR3022^12,13^ but with a different pose corresponding to a rotation of 73° about an axis perpendicular to the RBD α3 helix (central to both epitopes) (Fig. 2d,e). The Fab complex interface buries 564 and 361 Å^2^ of surface area for the CDRs of the heavy and light chains respectively. The EY6A interaction is mediated by the CDR loops H1, H2, H3, L1 and L3 contacting predominantly α3 but also α2 and the β2-α3, α4-α5 and α5-β4 loops of the RBD (Fig. 3 and Extended Data Fig. 6). A total of 16 residues from the heavy chain and 11 from the light chain participate in the interface together with 31 residues from the RBD. For the heavy chain these form potentially 6 hydrogen bonds and a single salt bridge between D99 (of H3) and K386 of the RBD and the light chain interface residues contribute an additional 6 hydrogen bonds. Hydrophobic interactions further increase the binding affinity (Fig. 3). Of the 31 residues involved in the interaction 21 are conserved between the CR3022 and EY6A epitopes (Fig. 3 and Extended Data Fig. 6). Conformational changes are introduced into the RBD by binding to EY6A at the α2 (residues 365-371) and α3 (residues 384-388) helices (Extended Data Fig. 6), similar to those seen for the CR3022 complex^12^. Comparison of the epitope residues for EY6A, CR3022^12^ and VHH-72^17^ shows that there is a very substantial overlap (Extended Data Fig. 6), although the bulk of the molecules extend in different directions, such that VHH-72 directly blocks ACE-2 binding^17^.

**Fig. 2.**
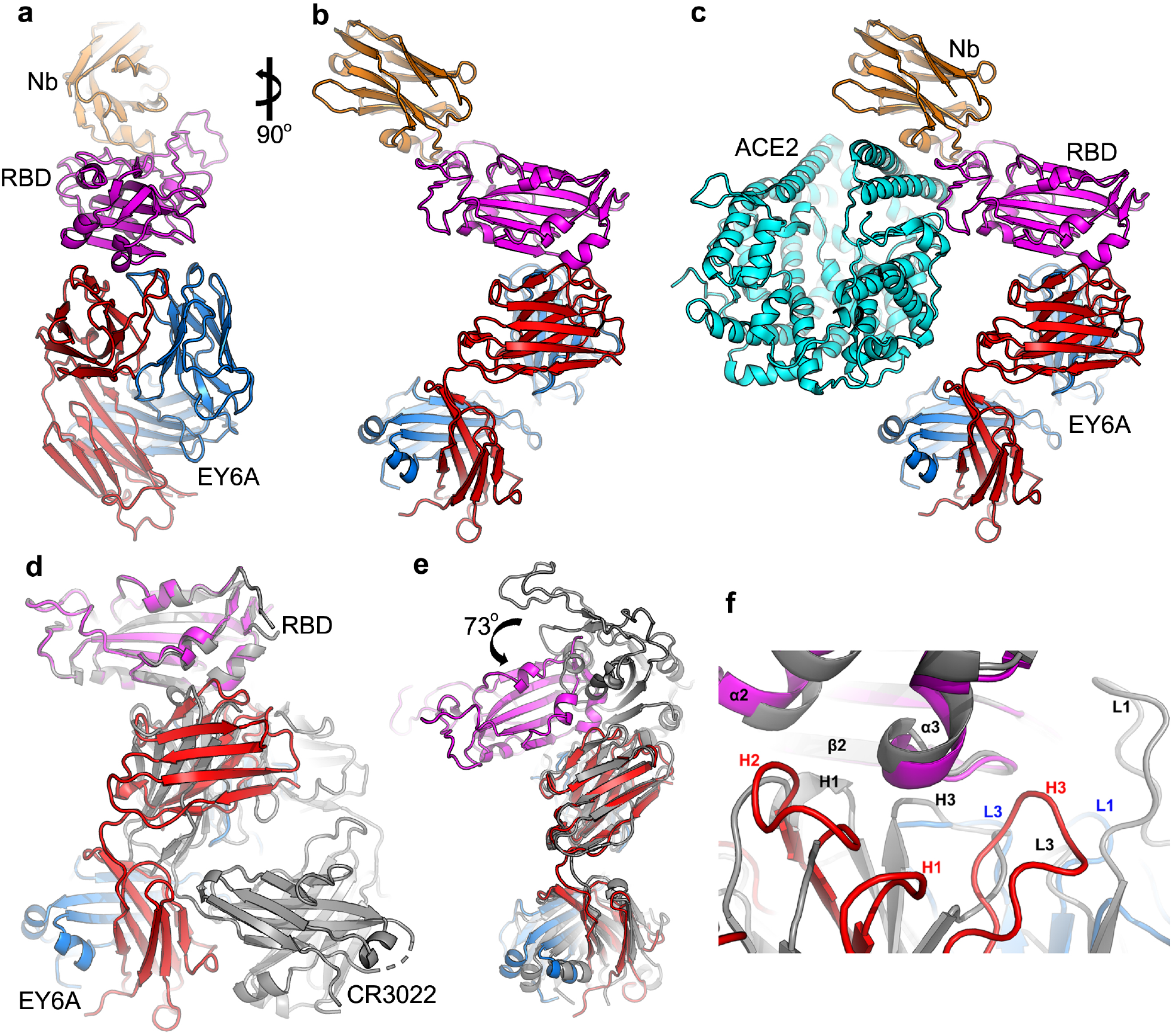
Overall structure of RBD-EY6A complex. **a**, A ribbon diagram of the crystal structure of RBD-EY6A-Nb ternary complex. The RBD, EY6A heavy and light chains, and Nb are coloured magenta, red, blue and orange, respectively. **b**, 90° rotation of (**a**). **c**, ACE2 (cyan) modelled into the ternary structure by superposing the RBD of the RBD-ACE2 complex (PDB ID, 6M0J) onto the ternary complex RBD. **d**, The RBD of the RBD-CR3022 complex (grey; PDB ID 6YLA) superposed on the RBD of the ternary complex, and **e**, as (**d**) but superposing the Vh domain. The Nb is omitted. **f**, Closeup of RBD-antibody interface of (**d**) showing the different epitope engagements by EY6A and CR3022.

**Fig. 3.**
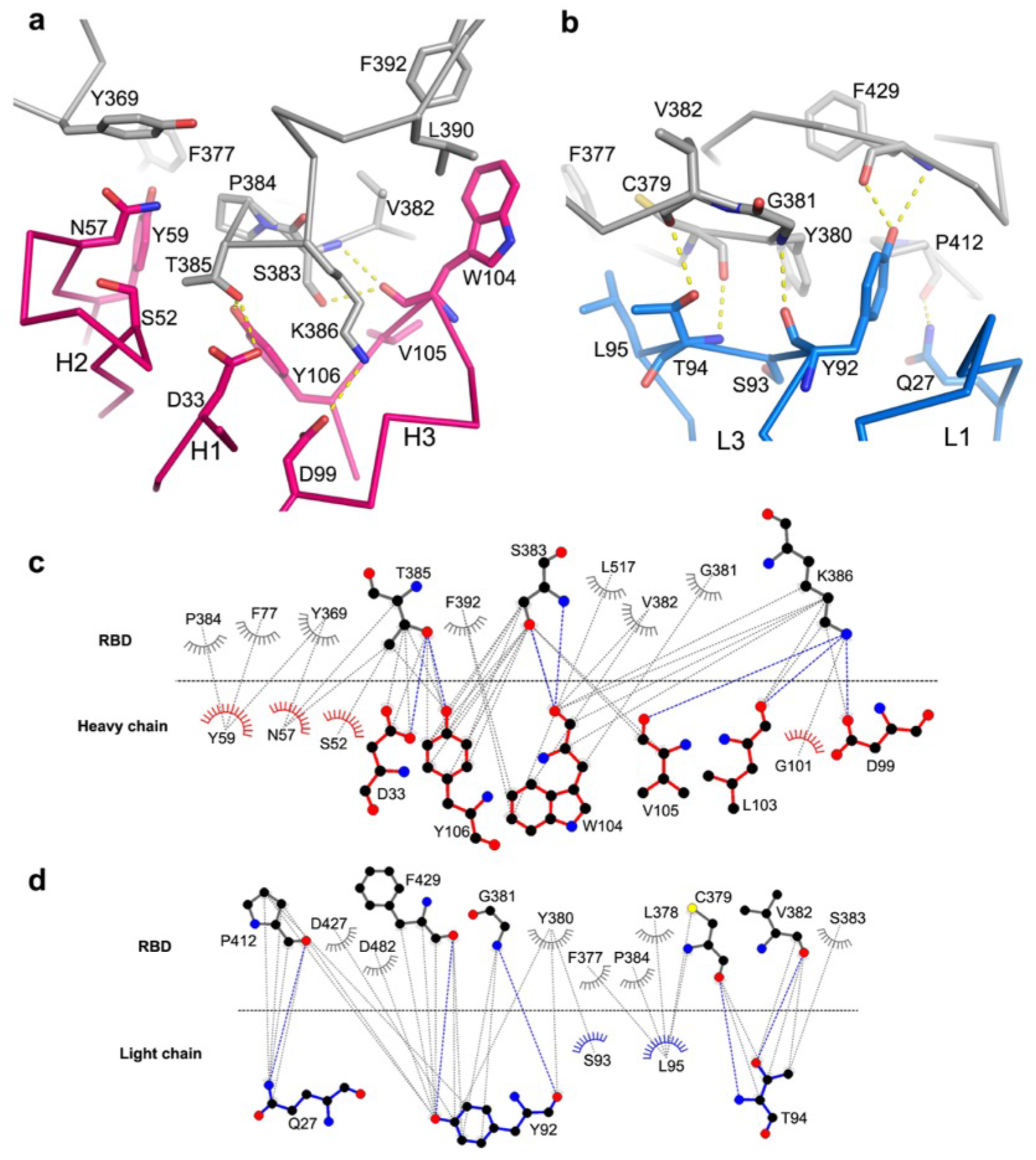
Details of RBD-EY6A interactions. **a**, Interactions between RBD and heavy chain, and **b**, between RBD and light chain. The RBD is shown in grey, heavy chain in red and light chain in blue. The side chains are drawn in thicker sticks and main chain Cα backbones in thinner sticks. Hydrogen bonds and salt bridges are shown as yellow broken sticks. **c**, LigPlot^44^ representation showing residues of the RBD involved in direct contacts with the EY6A heavy chain, and **d**, with the light chain. Potential hydrogen bonds ar shown with blue dotted lines.

### Residues that attach to EY6A form a protein-protein interface in the prefusion Spike

In the first pre-fusion Spike structures (PDB IDs: 6VSB^2^, 6VXX, 6VYB^4^), where residues 986 and 987 in the linker between two helices in S2 were mutated to a Pro-Pro sequence to prevent the conversion to the post-fusion helical conformation, the RBDs were found in either one ‘up’ two ‘down’ or all three ‘down’ configuration, and in both cases the epitope is inaccessible. In the ‘down’ position it is packed against another RBD of the trimer and the N-terminal domain (NTD) of the neighbouring protomer. A recent publication for the wild type Spike identifies a more closed form^18^ where the S1 portion of the Spike is tightened up. The structure is not yet deposited however, and so we have looked at the role of the epitope in the down rather than fully closed form, which will be broadly similar. Here the EY6A epitope packs down tightly against the S2 ‘knuckle’ bearing the Pro-Pro mutations, forming a buried protein-protein interface and making the epitope completely inaccessible. We assume that in the closed form this interaction will be even tighter and is probably responsible for maintaining the Spike in the pre-fusion state. Even when the RBD is in the ‘up’ configuration, the epitope remains largely inaccessible and substantial further movement of the RBD would be required to permit interaction unless more than one RBD was in the up conformation^12^.

### Cryo-EM analysis reveals that three EY6A Fabs can insert into the Spike

To investigate how the Fab insinuates itself into the Spike, we performed cryo-EM analysis. Spike ectodomain was mixed with a 6-fold molar excess of EY6A Fab and incubated at room temperature (21 °C) with an aliquot taken at 5 hours, applied to cryo-EM grids and frozen (Methods). Unbiased 2D class averages revealed three major particle classes with over one-third comprising a trimeric Spike/EY6A complex (some of which are self-associated) (Methods, Extended Data Table 4 and Figs. 6-10). Detailed analysis of this complex led to a reconstruction at 3.7 Å resolution (FSC = 0.143, C1 symmetry) which revealed three bound Fabs (Fig. 4a-d) nestled around the central axis at the top of the Spike. All three RBDs are in an ‘up and out’ configuration, markedly different to the published open forms (PDB:6VSB, 6VYB,^2,4^), being forced to rotate outwards by ~25°, such that the Spike is very open and appears on the verge of disruption (Fig. 4e.f). Indeed, the interactions with the Fab must partially stabilise what would otherwise be a disfavoured structure. This fragility is reflected in the observation that arrangement of RBD-EY6A complexes on the top of the Spike does not exactly follow 3-fold symmetry, with angles between the three RBDs being 120°, 119° and 121°. In addition, the orientations of the Vh domains relative to their associated RBDs differ slightly from that of the crystal structure (by 5°, 2° and 7°, respectively). The quality of the density suggests that these likely samples selected from a continuous distribution (Extended Data Fig. 10).

**Fig. 4.**
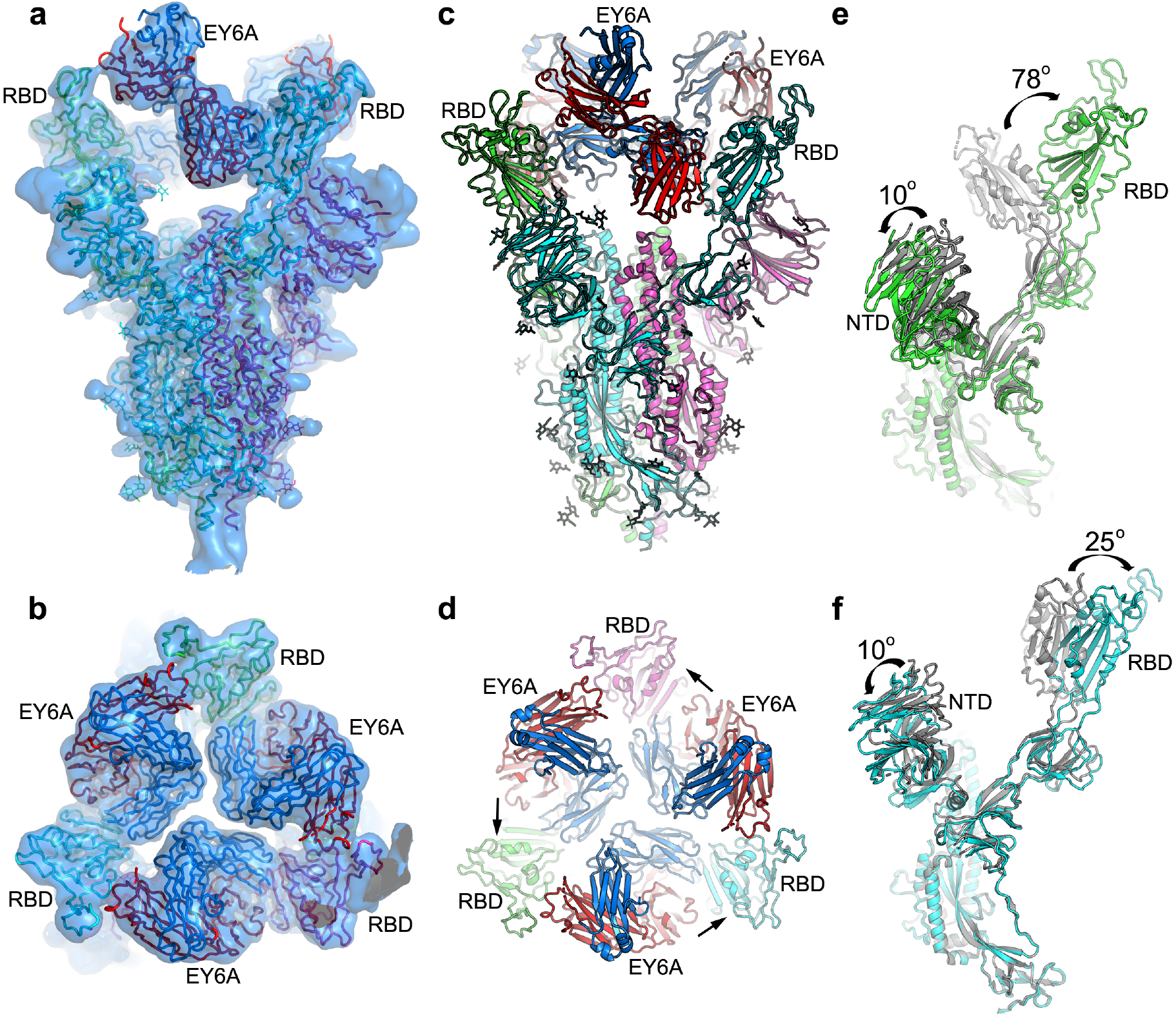
EM structure of SARS-CoV-2 spike and EY6A Fab complex. **a** & **b**, Side (**a**) and top (**b**) views of the cryo-EM density map drawn as semi-transparent surface showing three EY6A fabs bound to the Spike. **c** & **d**, Side and top views of the overall structure of the Spike-EY6A complex. Chains A, B and C of the Spike trimer and heavy and light chains of EY6A are shown in green, cyan, magenta, red and blue, respectively. The arrows in (**d**) indicate which RBD each Fab is binding. **e** & **f**, Comparison of the EY6A bound Spike structure with a reported open form Spike structure (grey; PDB ID 6VYB) with A chain in ‘down’ conformation (**e**) and B chain in ‘up’ conformation (**f**). NTD, N-terminal domain.

### Many Spikes lose structural integrity on EY6A incubation

The majority of the remaining particles form either a roughly 2-fold symmetric structure or a triangular association (Methods, Extended Data Table 4 and Figs. 7-11). Reconstructions of these particles were anisotropic due to a preferential orientation of the particles on the grid which was somewhat mitigated by collecting data with 30° tilt to yield reconstructions at 4.4 Å and 4.7 Å, respectively, in the plane of the grid but significantly worse resolution perpendicular to the grid (Extended Data Fig. 10). The reconstructions were sufficiently clear to allow the unambiguous fitting of EY6A-RBD complexes (Extended Data Fig. 11), however the density for what we assume are the N-terminal domains is poor in both reconstructions and we did not attempt to fit a model. These structures likely represent a residual well-structured fragment from the unfolding of the pre-fusion state of the Spike (SDS PAGE analysis shows that the Spike polypeptide remains largely uncleaved, Extended Data Fig. 12). The ‘dimeric’ and ‘trimeric’ structures are formed by different lateral associations and these also differ from that seen for similarly structurally degraded Spike-CR3022 complexes^12^ (Extended Data Fig. 11).

## Discussion

Convalescent serum has shown promise in patients severely ill with COVID-19^19,20^, thus immunotherapeutics have potential for treating COVID-19 even at a relatively late stage in the disease. To this end, it is desirable to find a combination of antibodies that neutralise the virus by different mechanisms to mitigate against immune evasion and antibody dependent enhancement. One neutralisation mechanism is blocking receptor attachment. We propose that attachment at the EY6A epitope is a further major neutralisation mechanism. In support of this, the epitope recognised by EY6A has been reported for several antibodies^12,13,21,22^ and nanobodies^17,23^ raised against SARS-CoV-2, SARS-CoV-1 and MERS. For SARS-CoV-1, CR3022 has also been shown to neutralise synergistically with ACE2 blocking antibodies^7^ Despite the spatial separation of the EY6A and ACE2 epitopes we find some cross-talk between the two binding events.

The EY6A epitope is extremely unusual, since it is completely inaccessible in the pre-fusion Spike trimer. This raises the question of what the mechanism of neutralisation might be. In the pre-fusion state the EY6A/CR3022 epitope rests down upon the upper end of the helixturn-helix between heptad repeat 1 (HR1) and the central helix (CH) of S2, essentially putting a lid on the spring-loaded extension of the helix which occurs on conversion to the postfusion state in the vicinity of the mutations designed to prevent conversion between the pre- and post-fusion conformation^24^ (Fig. 5). The residues of the epitope are crucial to these protein-protein interactions, and therefore highly conserved, explaining why it has, to date, proved impossible to generate mutations that escape binding of the antibody^7,12^. EY6A binding to the isolated RBD is tight (at ~2 nM it is roughly an order of magnitude tighter than CR3022) and, remarkably, the binding pose on top of the Spike allows three Fabs to bind simultaneously around the central axis (whereas CR3022 Fab cannot be accommodated). In line with this, a major portion of Spike molecules incubated for 5 h with EY6A are still in the intact pre-fusion state, with only about 1/3 being converted. Simple modelling suggests that a similar packing could occur for intact antibodies (Extended Data Fig. 13). In general, we would expect binders at this epitope to neutralise by displacing the ‘lid’ on the HR1/CH turn, reducing the stability of the pre-fusion state and therefore reducing the barrier to conversion to the more stable post-fusion trimer. This conversion is hindered in the construct we have used by the presence of the proline mutations at the turn between the helices. Premature conversion would prevent later attachment to the cell and block infectivity. The kinetics of this process will determine the effectiveness of the antibody in neutralisation and ultimately protection. Since the RBD is a relatively small domain there might also be an interplay between separate epitopes, thus we saw allosteric effects between EY6A and ACE2 binding and similarly VHH-72, which binds an overlapping epitope to EY6A, strongly inhibits ACE-2 binding by virtue of its different angle of attack^17^ The reason for the cross-talk between ACE2 and EY6A remains unclear, although we note that the glycan on ACE2 (residue N322) might well contribute to partial blocking (Extended Data Fig. 14). In summary attachment to this single epitope can cause neutralisation via more than one mechanism, and can exhibit strong synergy with ACE2 blocking antibodies^7^. Furthermore, due to the high level of conservation of key residues, tight binding antibodies targeting this epitope can neutralise a range of related viruses (usually spanning SARS-CoV-1 and SARS-CoV-2 and in some cases extending to MERS). We expect this remarkable epitope to be a major target for therapeutic exploitation.

**Fig. 5.**
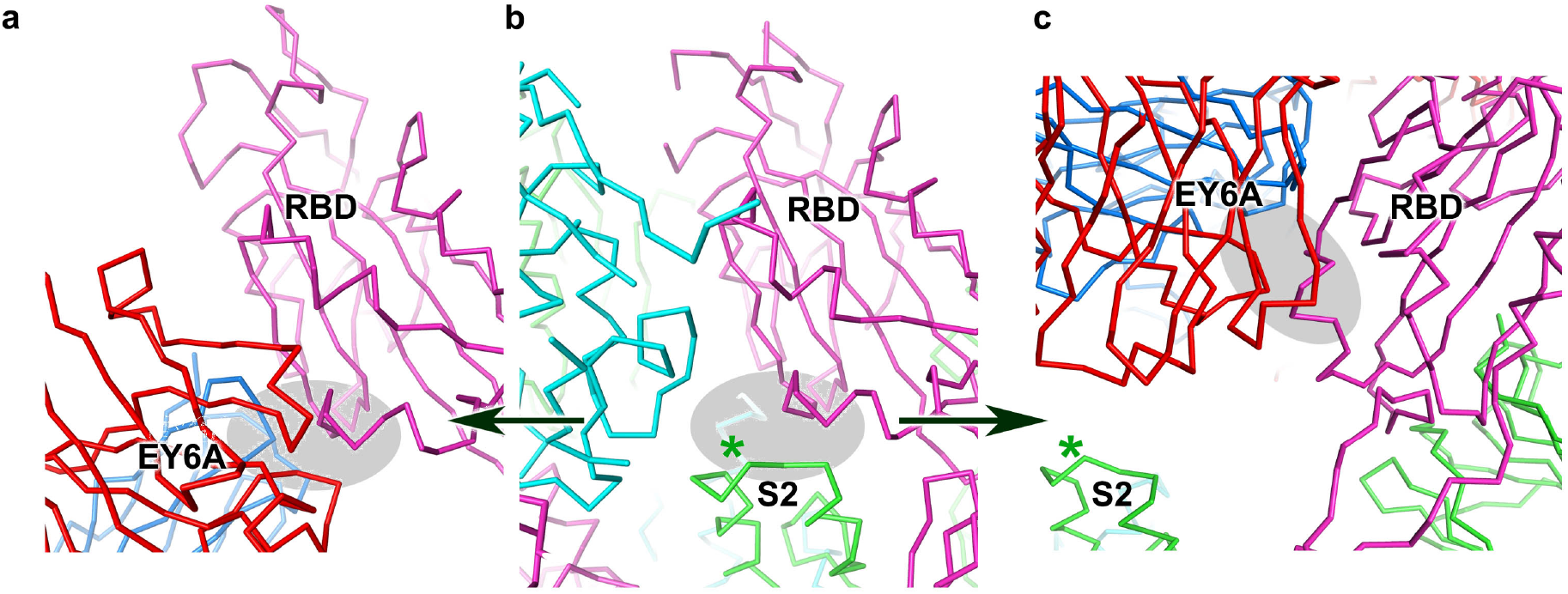
EY6A mimics S2 binding to the RBD. In panels **a** and **b** the orientation of the RBD is maintained demonstrating the commonality of the interaction area between EY6A and the S2 central helix. The star marks the position of the double proline mutations that prevent helix extension into the post-fusion state. Panel **c** maintains the orientation of S2 showing the displacement of the ‘lid’ from the S2 HR1/CH turn. The grey ellipse on all panels marks the region of the CR3022 epitope (the ‘lid’).

## Online Methods

### Antibody isolation study design

This study was designed to isolate SARS-CoV-2 antigen-specific human MAbs from peripheral plasmablasts in humans with natural SARS-CoV-2 infection, to characterize the antigenic specificity and phenotypic activity of SARS-CoV-2 Spike-reactive MAb, and to determine the structure of antibody in complex with viral antigen.

The infection of human by SARS-CoV-2 was confirmed by positive real-time reverse transcriptase polymerase chain reaction results of respiratory samples according to the guidelines of the Taiwan Centers for Disease Control (https://www.cdc.gov.tw/En). The study protocol and informed consent were approved by the ethics committee at the Chang Gung Medical Foundation and the Taoyuan General Hospital, Ministry of Health and Welfare, Taiwan. Each patient provided signed informed consent. The study and all associated methods were carried out in accordance with the approved protocol, the Declaration of Helsinki and Good Clinical Practice guidelines.

### Sorting of plasmablasts and production of human IgG monoclonal antibodies

Fresh peripheral blood mononuclear cells (PBMCs) were separated from whole blood by density gradient centrifugation and cryopreserved PBMCs were thawed. PBMCs were stained with a mix of fluorescent-labelled antibodies to cellular surface markers, including anti-CD3 (BD Biosciences, USA), anti-CD19 (BD Biosciences, USA), anti-CD27 (BD Biosciences, USA), anti-CD20 (BD Biosciences, USA), anti-CD38 (BD Biosciences, USA), anti-IgG (BD Biosciences, USA) and anti-IgM (BD Biosciences, USA). Plasmablasts were selected by gating on CD3-CD20-CD19+CD27hiCD38hiIgG+IgM-events and were isolated in chamber as single cell as previously described^26^. Sorted single cells were used to produce human IgG monoclonal antibodies as previously described^26^. Expression vectors that carry variable domains of heavy and light chains were transfected into the 293T cell line for expression of recombinant full-length human IgG monoclonal antibodies in serum-free transfection medium.

To determine the individual gene segments employed by VDJ and VJ rearrangements and the number of nucleotide mutations and amino acid replacements, the variable domain sequences were aligned with germline gene segments using the international ImMunoGeneTics (IMGT) alignment tool (http://www.imgt.org/IMGT_vquest/input).

### Protein cloning, expression and purification

#### EY6A IgG used for neutralisation and making Fab

Antibody was expressed using ExpiCHO expression system (LifeTechnologies) according to the manufacturer’s protocol and purified using Protein A MabSelect SuRE column (GE Healthcare). The wash buffer contained 20 mM Tris, 150 mM NaCl buffered to pH 8.6 and the elution was done using 0.1 M Citric acid pH 2.5. The eluate was neutralised immediately using 1.5 M Tris pH 8.6 and then buffer exchanged to phosphate-buffered saline (PBS) using 15 ml 30 kDa MWCO centrifugal filter (Merck Millipore).

#### Preparation of Fab-EY6A from IgG

EY6A Fab was digested from IgG with papain using a Pierce Fab Preparation Kit, following the manufacturer’s standard protocol.

#### Expression and purification of EY6A-6His Fab

Plasmids encoding the heavy and light chains of EY6A-6His Fab were amplified in Ecoli DH5α, then extracted and purified using a Qiagen HiSpeed Plasmid Giga Kit. HEK293T cells were transfected with the two plasmids. The medium was harvested and dialyzed into 1.7 mM NaH_2_PO_4_, 23 mM Na_2_HPO_4_, 250 mM NaCl, pH 8.0 at 4 °C overnight. The sample was then applied to a 5 ml HisTrap nickel column (GE Healthcare). Initially purified EY6A-6His Fab was then loaded onto a Superdex 75 HiLoad 16/60 gel filtration column (GE Healthcare) for further purification using 10 mM HEPES pH 7.4, 150 mM NaCl. Fractions containing EY6A-6His Fab were harvested and concentrated.

#### RBD, ACE2, Spike ectodomain and CR3022 cloning

Constructs are as described by Huo et al^12^.

#### Nanobody

This was derived from a naïve library followed by affinity maturation as described (Huo et al submitted).

#### Validation

All vectors were sequenced to confirm clones were correct.

#### Production of RBD and ACE2

These were transiently expressed in Expi293™ (Thermo Fisher Scientific, UK) and proteins were purified from culture supernatants by an immobilised metal affinity using an automated protocol implemented on an ÄKTAxpress (GE Healthcare, UK)^25^, followed by size-exclusion chromatography using a Hiload 16/60 superdex 75 or a Superdex 200 10/300GL column equilibrated in PBS pH 7.4 buffer. Recombinant Spike ectodomain was expressed by transient transfection in HEK293S GnTI-cells (ATCC CRL-3022) for 9 days at 30 °C. Conditioned media was dialysed against 2x phosphate buffered saline pH 7.4 buffer. The Spike ectodomain was purified by immobilised metal affinity chromatography using Talon resin (Takara Bio) charged with cobalt followed by size exclusion chromatography using HiLoad 16/60 Superdex 200 column in 150 mM NaCl, 10 mM HEPES pH 8.0, 0.02% NaN_3_ at 4 °C, before buffer exchange into 2 mM Tris pH 8.0, 200 mM NaCl^2^.

#### Deglycosylation of RBD

10 μL of Endoglycosidase F1 (~1mg/mL) was added to RBD (~2 mg/mL, 3 mL) and incubated at room temperature for two hours. RBD was then loaded to a Superdex 75 HiLoad 16/60 gel filtration column (GE Healthcare) for further purification using buffer 10 mM HEPES pH 7.4, 150 mM NaCl. Purified RBD was concentrated using a 10 kDa ultrafiltration tube (Amicon) to 12 mg/mL.

### Enzyme-linked immunosorbent assay (ELISA)

The ELISA plates (Corning^®^ 96-well Clear Polystyrene High Bind Stripwell™ Microplate, USA) were coated with SARS-CoV-2 antigen (Sino Biological, China) or SARS antigen (Sino Biological, China) or Middle East respiratory syndrome coronavirus (MERS) antigen (Sino Biological, China) or human coronavirus OC43 antigen (Sino Biological, China) at optimal concentration in carbonate buffer and incubated at 4°C overnight. The next day unbound antigens were removed by pipetting to avoid risk of forming aerosols. Nonspecific binding was blocked with the solution of phosphate-buffered saline with 3% bovine serum albumin at room temperature for 1 hour on a shaker. After removing blocking buffer, monoclonal antibody-containing cell culture supernatant or purified monoclonal antibody preparation were added and incubated at 37°C for 1 hour. The non-transfected cell culture supernatant, anti-influenza human monoclonal antibody BS 1A (in house), anti-SARS Spike monoclonal antibody CR3022 and convalescent serum were used as antibody controls for each experiment. After incubation, the plate was washed and incubated with horseradish peroxidase-conjugated rabbit anti-human IgG (Rockland Immunochemicals, USA) as secondary antibody. After incubation, the plate was washed and developed with TMB substrate reagent (BD Biosciences, USA). Reaction was stopped by 0.5M Hydrochloric acid and the optical density was measured at OD450 on a microplate reader. The well that yielded an OD value four times the mean absorbance of negative controls (BS 1A) was considered positive.

### Immunofluorescence assay

SARS-CoV-2 (strain CDC-4) infected Vero E6 cells were prepared and fixed with acetone in the biosafety level 3 (BSL-3) laboratory following biosafety rules and guidelines^27^. The fixed cells on the cover slips were stained with monoclonal antibody-containing cell culture supernatant. The anti-influenza human monoclonal antibody BS 1A (in house), anti-SARS Spike monoclonal antibody CR3022 and convalescent serum were used as controls. Following incubation and wash, the cells were stained with FITC-conjugated goat anti-human IgG secondary antibody (Invitrogen, USA) and binding antibodies were detected by fluorescence microscopy.

### Quantitative PCR-based neutralisation assay

Neutralization activity of monoclonal antibody-containing supernatant was measured using a SARS-CoV-2 (strain CDC-4) infection of Vero E6 cells^27^. Briefly, Vero E6 cells were preseeded in a 96 well plate at a concentration of 2 x 10^4^ cells per well. On the following day, monoclonal antibody-containing supernatant were mixed with an equal volume of 100 TCID50 virus preparation and incubated at 37 °C for 1 hour. The mixture was added into seeded Vero E6 cells and incubated at 37 °C for 5 days. The cell control, virus control, and virus back-titration were setup for each experiment. At day 5, the culture supernatant was harvested from each well and the viral RNA was extracted by the automatic LabTurbo system (Taigen, Taiwan) following the manufacturer’s instructions. for the most part, except that the specimen was pretreated with Proteinase K prior to RNA extraction. Real-time reverse transcription polymerase chain reaction was performed in a 25-μL reaction containing 5 μL of RNA^28^. The primers and probe used to amplify the E gene were as follows: E_Sarbeco_F: 5’-ACAGGTACGTTAATAGTTAATAGCGT-3’, E_Sarbeco_R: 5’-ATATTGCAGCAGTACGCACACA-3’, E_Sarbeco_P1: FAM-ACACTAGCCATCCTTACTGCGCTTCG-BBQ.

### Cell-based ACE2 blocking assays

MDCK-SIAT1 cells were stably transfected with human ACE2 cDNA using a second-generation lentiviral vector. ACE2 expressing cells were then FACS sorted post staining with RBD-6H followed by a secondary anti-His AlexaFluor 647 labelled antibody. Cells (3 x 10^4^ per well) were seeded the day before the assay in a flat-bottomed 96-well plates. A serial half-log dilution (ranging 1 μM to 0.1 nM) of antibodies and controls were performed in 30 μL volume. PBS supplemented with 0.1 % BSA (37525; Thermo Fisher Scientific) was used for dilution of all antibodies. 30 μL of biotinylated RBD (using EZ-link Sulfo-NHS-Biotin (A39256; Life Technologies)) at 25 nM was added to the titrated antibodies. Cells were washed with PBS and 50 μL of each mixture of biotinylated RBD and an antibody was transferred to the MDCK-ACE2 and incubated for 1 h at room temperature. Cells were then washed with PBS and incubated for 1 h with a second layer Streptavidin-HRP antibody (434323, Life Technologies) diluted to 1:1,600 and developed with BM POD Substrate (11484281001, Roche) for 5 min before stopping with 1M H_2_SO_4_. Plates were then read on a ClarioStar Plate Reader and graphs were plotted as % binding of biotinylated ACE2 to RBD. Binding % = (X - Min)/ (Max - Min)*100 where X = Measurement of the competing component, Min = Buffer without binder biotinylated RBD, Max = Biotinylated RBD alone. Inhibitory concentration at 50 % (IC_50_) of the antibodies against RBD was determined using non-linear regression [inhibitor] versus normalised response curve fit using GraphPad Prism 8.

### Surface plasmon resonance

Surface plasmon resonance experiments were performed using a Biacore T200 (GE Healthcare). All assays were performed using a Sensor Chip Protein A (GE Healthcare), with a running buffer of PBS pH 7.4 supplemented with 0.005% v/v Surfactant P20 (GE Healthcare) at 25 °C. To determine the binding kinetics between the RBD of SARS-CoV-2 and EY6A mAb, two different experimental settings were attempted. The first experiment was performed with RBD-Fc immobilized on the sample flow cell. The reference flow cell was left blank. The EY6A Fab was injected over the two flow cells at a range of five concentrations prepared by serial two-fold dilution from 50 nM, at a flow rate of 30 μL/min using a Single-cycle kinetics program with an association time of 75 s and a dissociation time of 900 s. Running buffer was also injected using the same program for background subtraction. The second experiment was performed with EY6A IgG immobilised on the sample flow cell. The reference flow cell was left blank. The RBD was injected over the two flow cells at a range of five concentrations prepared by serial two-fold dilution from 100 nM, at a flow rate of 30 μL/min using a Single-cycle kinetics program with an association time of 90 s and a dissociation time of 60 s. Running buffer was also injected using the same program for background subtraction. All data were fitted to a 1:1 binding model using the Biacore T200 Evaluation Software 3.1. In the competition assay where CR3022 IgG or ACE2-hIgG1Fc was used as the ligand, the following samples were injected: (1) a mixture of 1 μM EY6A Fab and 0.1 μM RBD, (2) a mixture of 1 μM (anti-Caspr2) E08R Fab and 0.1 μM RBD, (3) 0.1 μM RBD, (4) 1 μM EY6A Fab; (5) 1 μM E08R Fab. In the competition assay where EY6A IgG was used as the ligand, the following samples were injected: (1) a mixture of 1 μM CR3022 Fab and 0.1 μM RBD, (2) a mixture of 1 μM ACE2 and 0.1 μM RBD, (3) a mixture of 1 μM E08R Fab and 0.1 μM RBD, (4) 0.1 μM RBD, (5) 1 μM CR3022 Fab, (6) 1 μM ACE2, (7) 1 μM E08R Fab. All injections were performed with an association time of 60 s and a dissociation time of 600 s. All curves were plotted using GraphPad Prism 8.

### Plaque reduction neutralisation test (PHE, Porton Down)

SARS-CoV-2 (Australia/VIC01/2020)^29^ was diluted to a concentration of 933 pfu ml^-1^ (70 pfu/75 μl) and mixed 50:50 in minimal essential medium (MEM) (Life Technologies, California, USA) containing 1 % foetal bovine serum (Life Technologies) and 25 mM HEPES buffer (Sigma, Dorset, UK) with doubling antibody dilutions in a 96-well V-bottomed plate. The plate was incubated at 37°C in a humidified box for 1 h to allow neutralisation to take place. before the virus-antibody mixture was transferred into the wells of a twice Dulbecco’s PBS-washed 24-well plate containing confluent monolayers of Vero E6 cells (ECACC 85020206; PHE, UK) that had been cultured in MEM containing 10 % (v/v) FBS. Virus was allowed to adsorb onto cells at 37°C for a further hour in a humidified box, and overlaid with MEM containing 1.5 % carboxymethylcellulose (Sigma), 4 % (v/v) FBS and 25mM HEPES buffer. After 5 days incubation at 37°C in a humidified box, the plates were fixed overnight with 20 % formalin/PBS (v/v), washed with tap water and then stained with 0.2 % crystal violet solution (Sigma) and plaques were counted. Median neutralising titres (ND_50_) were determined using the Spearman-Karber formula^30^ relative to virus only control wells.

### Plaque reduction neutralisation test (Oxford)

Plaque reduction neutralization tests were performed using passage 4 of SARS-CoV-2 Victoria/01/2020^29^. Virus suspension at appropriate concentrations in Dulbecco’s Modification of Eagle’s Medium containing 1 % FBS (D1; 100 μL) was mixed antibody (100 μL) diluted in D1 at a final concentration of 50 μg/mL, 25 μg/mL, 12.5 ug/mL or 6.125 μg/mL, in triplicate, in wells of a 24 well tissue culture plate, and incubated at room temperature for 30 minutes. Thereafter, 0.5 mL of a single cell suspension of Vero E6 cells in D1 at 5 x 105/mL was added, and incubated for 2 h at 37°C before being overlain with 0.5 mL of D1 supplemented with carboxymethyl cellulose (1.5 %). Cultures were incubated for a further 4 days at 37°C before plaques were revealed by staining the cell monolayers with amido black in acetic acid/methanol.

### Crystallisation, data collection and X-ray structure determination

Purified and deglycosylated RBD and EY6A Fab were combined in an approximate molar ratio of 1:1 at a concentration of 6.5 mg/mL. Nb was also combined with EY6A-6His Fab and RBD in a 1:1:1 molar ratio with a final concentration of 5.7 mg/mL. These two complexes were separately incubated at room temperature for one hour. Initial screening of crystals was performed in Crystalquick 96-well X plates (Greiner Bio-One) with a Cartesian Robot using the nanolitre sitting-drop vapour diffusion method as previously described^31,32^. Crystals for the binary complex were initially obtained from Hampton Research Index screen, condition B7 containing 0.04 M NaH_2_PO_4_, 0.96 M K_2_HPO_4_ and further optimized to produce better crystals in 0.02 M NaH_2_PO_4_, 0.98 M K_2_HPO_4_. Good crystals for the ternary complex were also obtained from the Index screen condition G1 containing 25 % (w/v) PEG 3350, 0.2 M NaCl, 0.1 M Tris pH 8.5.

Crystals were soaked in a solution containing 25% glycerol and 75% reservoir solution for a few seconds and then mounted in loops and frozen in liquid nitrogen prior to data collection. Diffraction data were collected at 100 K at beamline I03 of Diamond Light Source, UK. Diffraction images of 0.1° rotation were recorded on an Eiger2 XE 16M detector with an exposure time of 0.01 s per frame, beam size 80×20 μm and 100% beam transmission. Data were indexed, integrated and scaled with the automated data processing program Xia2-dials^33,34^ The data set for the binary complex of 360° was collected from a single frozen crystal to 3.8 Å resolution with 20-fold redundancy. The crystal belongs to space group *P3_1_21* with unit cell dimensions *a* = *b* = 166.6 Å and *c* = 270.8 Å. The structure was determined by molecular replacement with PHASER^35^ using search models of antibody CR3022 Fab and the RBD of the RBD/CR3022 Fab complex (PDB ID 6YLA;^12^). There are three RBD/EY6A complexes in the crystal asymmetric unit, resulting in a crystal solvent content of ~75%. For the ternary complex, a data set of 360° rotation with data extending to 2.6 Å was collected on beamline I03 of Diamond with exposure time 0.008 s per 0.1° frame, beam size 80×20 μm and 100% beam transmission). The crystal also belongs to space group *R3* but with unit cell dimensions (*a* = *b* = 178.1 Å and *c* = 87.8 Å). There is one RBD/EY6A/Nb complex in the asymmetric unit and a solvent content of ~61%.

### X-ray crystallographic refinement and electron density map generation

One cycle of REFMAC5^36^ was used to refine atomic coordinates after manual correction in COOT^37^ to the protein sequence from the search model. For both the binary and ternary complexes the final refinement used PHENIX^38^ resulting in R_work_ = 0.219 and R_free_ = 0.259 for all data to 3.8 Å resolution for the binary complex and to R_work_ = 0.216 and R_free_ = 0.262 for all data to 2.64 Å resolution for the ternary complex. There is well ordered density for a single glycan at the glycosylation site N343 in the RBD.

Data collection and structure refinement statistics are given in Extended Data Table 3. Structural comparisons used SHP^39^, residues forming the RBD/Fab interface were identified with PISA^40^, figures were prepared with PyMOL (The PyMOL Molecular Graphics System, Version 1.2r3pre, Schrödinger, LLC).

### EY6A Fab / Spike complex preparation and cryo-EM data collection

Spike protein, following SEC purification, was buffer exchanged into 2 mM Tris pH 8.0, 200 mM NaCl, 0.02 % NaN_3_ buffer using a desalting column (Zeba, Thermo Fisher). A final concentration of 0.18 mg/mL was incubated with EY6A Fab (in the same buffer) in a 6:1 molar ratio (Fab to trimeric Spike) at room temperature for 5 hrs. Control grids of Spike alone after incubation at room temperature for 5 hrs were also prepared.

Each grid was prepared using 3 μL sample applied to a freshly glow-discharged on high for 20 s (Plasma Cleaner PDC-002-CE, Harrick Plasma) holey carbon-coated 200mesh copper grid (C-Flat, CF-2/1, Protochips) and excess liquid removed by blotting for 5-5.5 s with a blotting force of −1 using Vitrobot filter paper (grade 595, Ted Pella Inc.) at 4.5 °C, 100 % relative humidity and immediately plunge frozen into ethane slush using a Vitrobot Mark IV (Thermo Fisher).

Grids were screened on a Titan Krios microscope using SerialEM operating at 300 kV (Thermo Fisher). Movies were collected on a K3 detector on a Titan Krios operating at 300 kV in super resolution mode, with a calibrated Super Resolution pixel size of 0.415 A/pix at both 0° and 30° tilt. To compensate for the poorer contrast with tilted data, it was necessary to use a higher dose rate for the latter dataset.

### Cryo-EM Data Processing

Alignment and motion correction was performed using Relion3.1’s implementation of motion correction^41^, with a 5-by-5 patch-based alignment. All frames were binned by two, resulting in a final calibrated pixel size of 0.83 Å/pixel. Contrast-transfer-function (CTF) of full-dose and non-weighted micrographs was estimated within a CryoSPARC wrapper for Gctf-v1.06^42^. Images were then manually inspected and those with poor CTF-fits were discarded. Particles were then picked by unbiased blob picking in CryoSPARC v.2.14.1^43^ and subjected to rounds of 2D classification.

For the Spike-EY6A dataset (structure A), 2,096,246 Spike-like particles were used to make a template to pick particles from the untilted dataset, which were then filtered by 2D classification to 110,096 particles and then further refined by 3D classification with an ab initio model set. For the 30 ° dataset, 124,194 particles were used as a template, and filtered by 2D classification to a set of 84,230 particles and then, as before, further refined by unbiased 3D classification. The two particle sets were then refined together, with a final set of 144,680 particles.

For B and C (triangular ring and ‘dimeric’ form), particles from both the zero and 30° datasets were combined in a similar manner to the Spike-EY6A dataset using the ‘Exposure Group Utilities’ module in CryoSPARC. Both particle sets (B, 41372 particles and C, 119,343 particles) were then reclassified and the best class refined with non-uniform refinement. For B, C3 symmetry was imposed at this final refinement stage, resulting in an appreciable improvement in resolution, as indicated by inspection and gold-standard FSC = 0.143 (4.7 versus 5.9 Å, see Extended Data Table 4).

### Cryo-EM model building and refinement

The EM density of Spike/EY6A was fitted with the structure of a closed form of Spike (PDB ID 6VXX) apart from the RBDs and EY6A Fab which were fitted with RBD/EY6A of the ternary crystal structure using COOT^37^. Due to the lower resolution, RBD and EY6A are only fitted to the ‘dimeric’ and ‘trimeric’ EM density. The Spike/EY6A structure was refined with PHENIX^38^ real space refinement, first as a rigid body and then by positional and B-factor refinements. Only rigid body refinement was applied to the ‘dimeric’ and the ‘trimeric’ complexes. The statistics of EM data collection and structure refinement are shown in Extended Data Table 4.

## Acknowledgements

We acknowledge the BD FACSAria™ cell sorter service provided by the Core Instrument Center of Chang Gung University. The works of sorting of plasmablasts, production and characterization of human MAbs were supported by the Chang Gung Memorial Hospital (BMRPE22). This work was supported by the Chinese Academy of Medical Sciences (CAMS) Innovation Fund for Medical Science (CIFMS), China (grant number: 2018-I2M-2-002) to D.I.S. E.E.F, H.M.E.D. and J.Ren are supported by the Wellcome Trust (101122/Z/13/Z), Y.Z. by Cancer Research UK (C375/A17721) and D.I.S. and E.E.F. by the UK Medical Research Council (MR/N00065X/1). J.H. is supported by a grant from the EPA Cephalosporin Fund. PPUK is funded by the Rosalind Franklin Institute EPSRC Grant no. EP/S025243/1. The National Institute for Health Research Biomedical Research Centre Funding Scheme supports G.R.S. together with the Chinese Academy of Medical Sciences (CAMS) Innovation Fund for Medical Science (CIFMS), China (grant number: 2018-I2M-2-002), which also supports D.I.S. We are grateful also grateful for a Fast Grant from Fast Grants, Mercatus Center to support the isolation of human monoclonal antibodies to SARS-2. G.R.S. is also supported as a Wellcome Trust Senior Investigator (grant 095541/A/11/Z). P.R and A.R.M are funded by the T.K.T is funded by the EPA Cephalosporin Early Career Teaching and Research Fellowship and the Townsend-Jeantet Charitable Trust (Charity Number 1011770). T.M. is supported by Cancer Research UK grants C20724/A14414 and C20724/A26752 to Christian Siebold. This is a contribution from the UK Instruct-ERIC Centre. The Wellcome Centre for Human Genetics is supported by the Wellcome Trust (grant 090532/Z/09/Z). Virus used for the neutralisation assays was a gift from Julian Druce, Doherty Centre, Melbourne, Australia. We acknowledge Diamond Light Source for time on Beamline I03 under Proposal mx19946 and for electron microscope time at the UK national electron bio-imaging centre (eBIC), Proposal BI26983, both COVID-19 Rapid Access. D.I.S. is a Jenner Investigator. This is a contribution form the UK Instruct-ERIC Centre. Huge thanks to the teams, especially at the Diamond Light Source and Department of Structural Biology, Oxford University that have enabled work to continue during the pandemic.

## Author Information

These authors contributed equally: D.Z., HMED, C.-P.C.

## Contributions

J.H. performed interaction analyses and T.K.T., P.R., R.F.D., A.R.T., K.B., K.G., R.F.D. and M.C. prepared material for, and executed, neutralisation assays and the cell-based ACE2 blocking assays. Y.Z., D.Z., J.H., J.Ren, performed sample preparation for and crystallographic experiments and processed the data. N.G.P. assisted with X-ray diffraction data collection. J.Ren refined the structures and together with E.E.F. and D.I.S. analysed the results. G.R.S., J.M., and P.S. prepared the Spike construct. L.C. helped performed cryo-EM data processing, T.M. prepared the Spike sample, H.M.E.D. R.R.R. and P.N.M.S. prepared cryo-EM grids, H.M.E.D. performed cryo-EM sample preparation, screening and processing and J.Raedecke performed cryo-EM data collection, and J.Ren refined the cryo-EM structures. E.E.F., J.Ren, Y.Z. K.-Y.A.H. and D.I.S. wrote the manuscript. K.-Y.A.H. isolated and characterized EY6A. C.-P.C., C.-G.H., T.-H.C., S.-R.S, Y.-C.L., C.-Y.C., S.-H.C., Y.-C.H., T.-Y.L., and C.M helped prepare materials, perform experiments and analysed data. All authors read and approved the manuscript.

## Corresponding authors

Correspondence to David I. Stuart or Kuan-Ying A. Huang.

## Data availability

The coordinates and structure factors of the SARS-CoV-2 RBD/EY6A crystallographic complexes are available from the PDB with accession codes XXX and VVV respectively. EM maps and structure models are deposited in EMDB and PDB with accession codes XXX and YYY for the pre-fusion Spike, and XXXXX and yyyy for the dimeric complex respectively. The data that support the findings of this study are available from the corresponding authors on request.

