## Extended Data for "Structural basis for the neutralization of SARS-CoV-2 by an antibody from a convalescent patient"

Figs 1-14.

Tables 1- 4.

**a** **b**

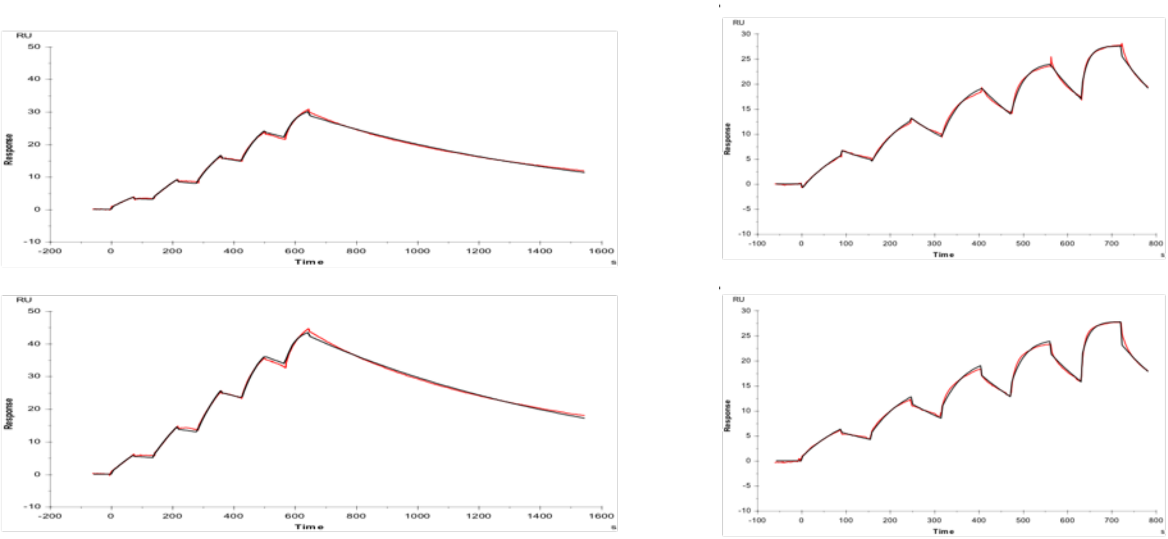


**Extended Data Fig. 1 | Binding affinity between RBD and EY6A Fab. a-b,** Surface plasmon resonance binding sensorgrams. **a,** RBD-Fc was immobilised as the ligand and EY6A Fab was used as analyte at five concentrations (3.125, 6.25, 12.5, 25 and 50 nM). **b,** EY6A IgG was immobilised as the ligand and RBD was used as analyte at five concentrations (6.25, 12.5, 25 50 and 100 nM). The average kinetic values from these two sets of experiment are listed in Extended Data Table 1.

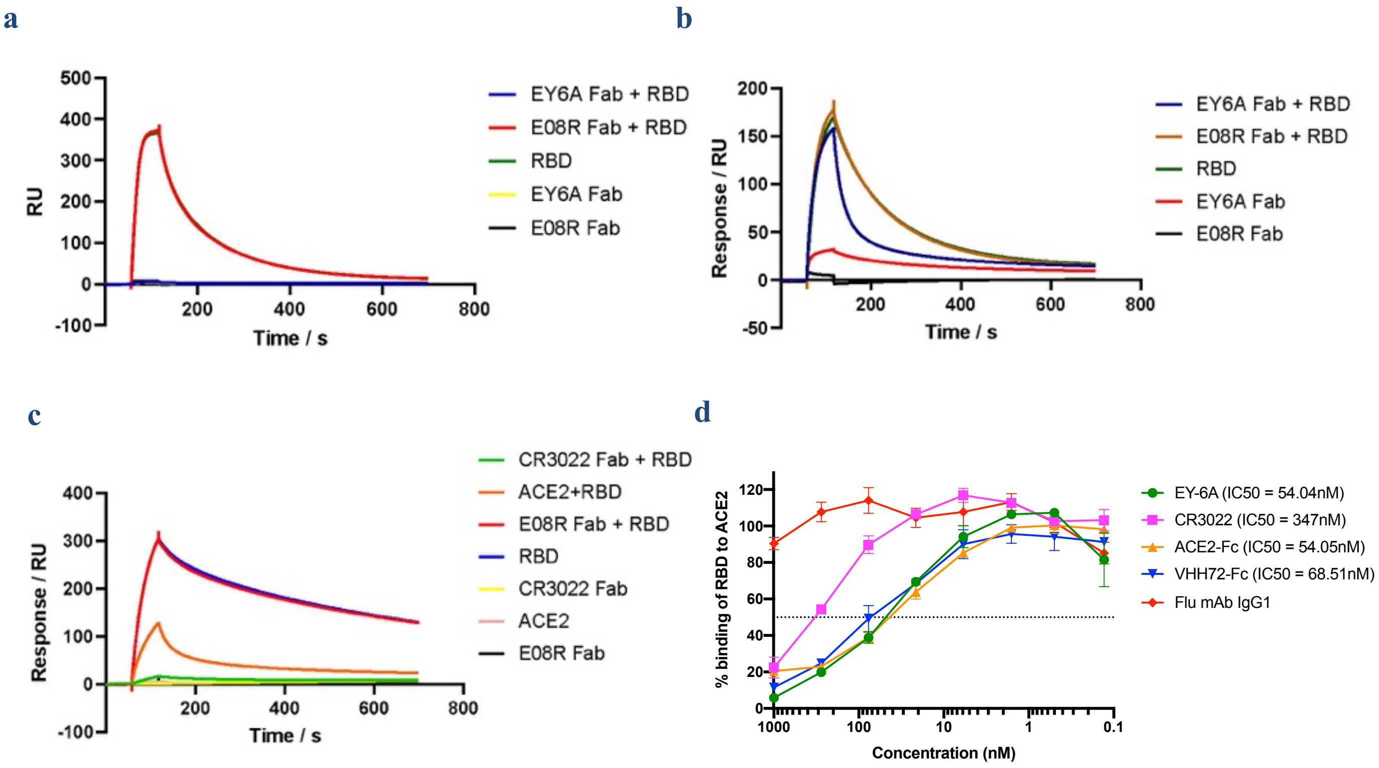


**Extended Data Fig. 2 | Binding competition of EY6A and ACE2 for RBD.** Surface plasmon resonance binding sensorgrams. **a,** CR3022 IgG was immobilised, the following samples were injected: (1) a mixture of 1 μM EY6A Fab and 0.1 μM RBD; (2) a mixture of 1 μM E08R Fab and 0.1 μM RBD; (3) 0.1 μM RBD; (4) 1 μM EY6A Fab; (5) 1 μM E08R Fab. **b,** ACE2-hIgG1Fc was immobilised, the following samples were injected: (1) a mixture of 1 μM EY6A Fab and 0.1 μM RBD; (2) a mixture of 1 μM E08R Fab and 0.1 μM RBD; (3) 0.1 μM RBD; (4) 1 μM EY6A Fab; (5) 1 μM E08R Fab. **c,** EY6A IgG was immobilised, the following samples were injected: (1) a mixture of 1 μM CR3022 Fab and 0.1 μM RBD, (2) a mixture of 1 μM ACE2 and 0.1 μM RBD, (3) a mixture of 1 μM E08R Fab and 0.1 μM RBD, (4) 0.1 μM RBD, (5) 1 μM CR3022 Fab, (6) 1 μM ACE2, (7) 1 μM E08R Fab. **d,** Purified antibodies, ACE2-Fc or VHH72-Fc were added at various concentrations to biotin labelled RBD-6H (25nM) and added to MDCK-SIAT1 cells stably expressing human ACE2 on the cell surface (MDCK-ACE2). The amount of biotinylated RBD bound to the cell was measured.

Experiments were performed in duplicate with the mean +/- SD are shown. $IC_{50}$ is calculated by linear interpolation as described in Methods.

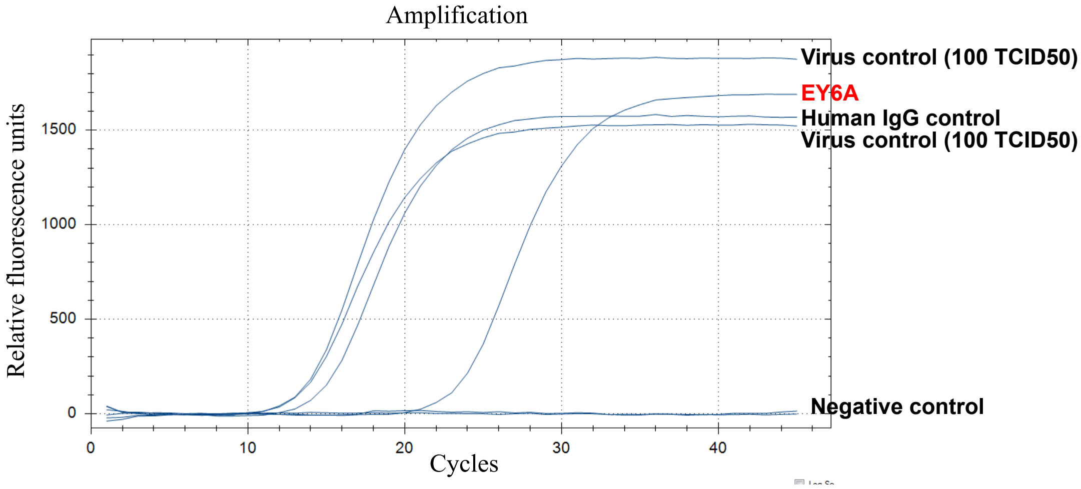


| Ct value of culture supernatant, after 5 days of incubation | |
|---|---|
| **Virus control supernatant (100 TCID50)** | **13.0** |
| **EY6A supernatant (1.5 µg/ml)** | **22.7*** |
| **Human IgG control supernatant (1.5 µg/ml)** | **13.1** |

**Extended Data Fig. 3 | Neutralisation of SARS-CoV-2 by EY6A.** Neutralisation Data by measuring Ct value of virus signal in the supernatant of SARS-CoV-2 infected Vero E6 cells in an E gene-based real-time reverse-transcription PCR assay [27]. An increase indicates a decrease in virus template. Each unit increase suggests a 2x reduction resulting from presence of Mab. A ~10x increase in Ct = ~1,000 fold reduction of virus. Anti-influenza H3 MAb BS 1A was included as a human IgG control in the assay. The neutralization assay was carried out twice with equivalent results. * = produced in bulk.

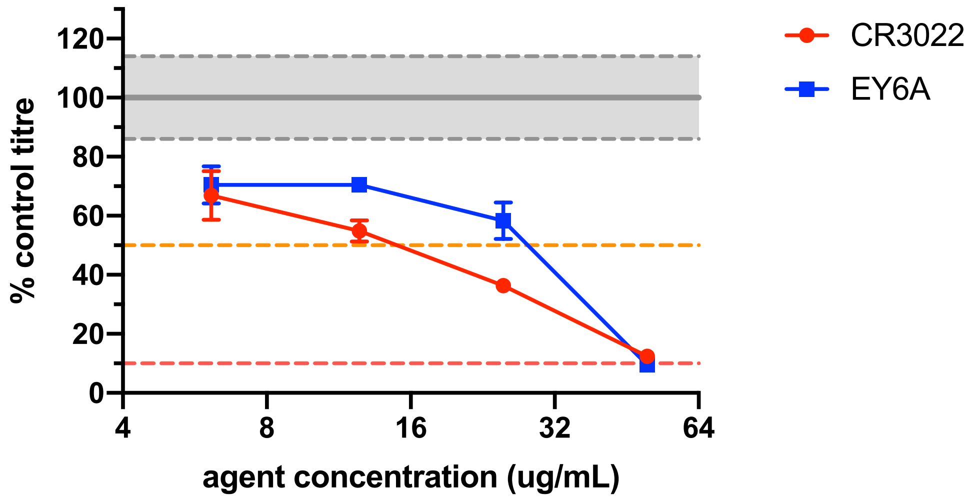


**Extended Data Fig. 4 | Neutralisation of SARS-CoV-2 by EY6A and CR3022.** Oxford Vero cell based PRNT assay.

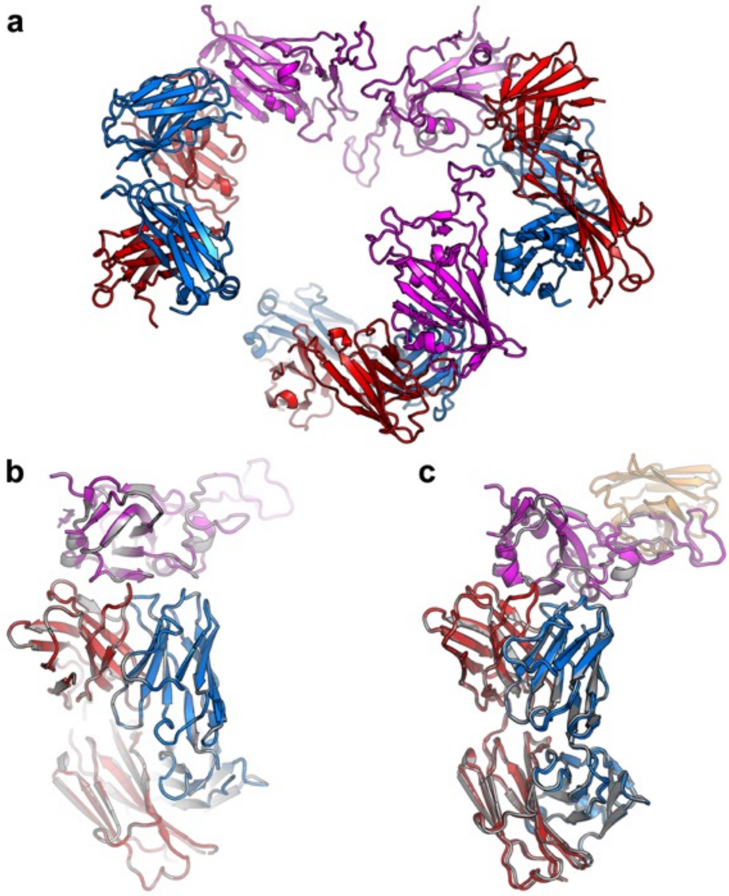


**Extended Data Fig. 5 | Comparison of the binding poses in crystal structures of the binary and ternary complexes. a**, Three RBD-EY6A binary complexes in the crystal asymmetric unit. RBD is shown in magenta, EY6A heavy chain in red and light chain in blue. **b**, Superimposition of the three RBD-EY6A complexes in the asymmetric unit showing the same binding pose. One complex is shown in colour as in (**a**) and the other two in grey. **c**, Comparison of RBD-EY6A binary complex with RBD-EY6A-Nb ternary complex by overlapping the RBDs, the RBD and EY6A are shown in colour as in (**a**) with Nb in orange in the ternary complex, and grey in the binary complex.

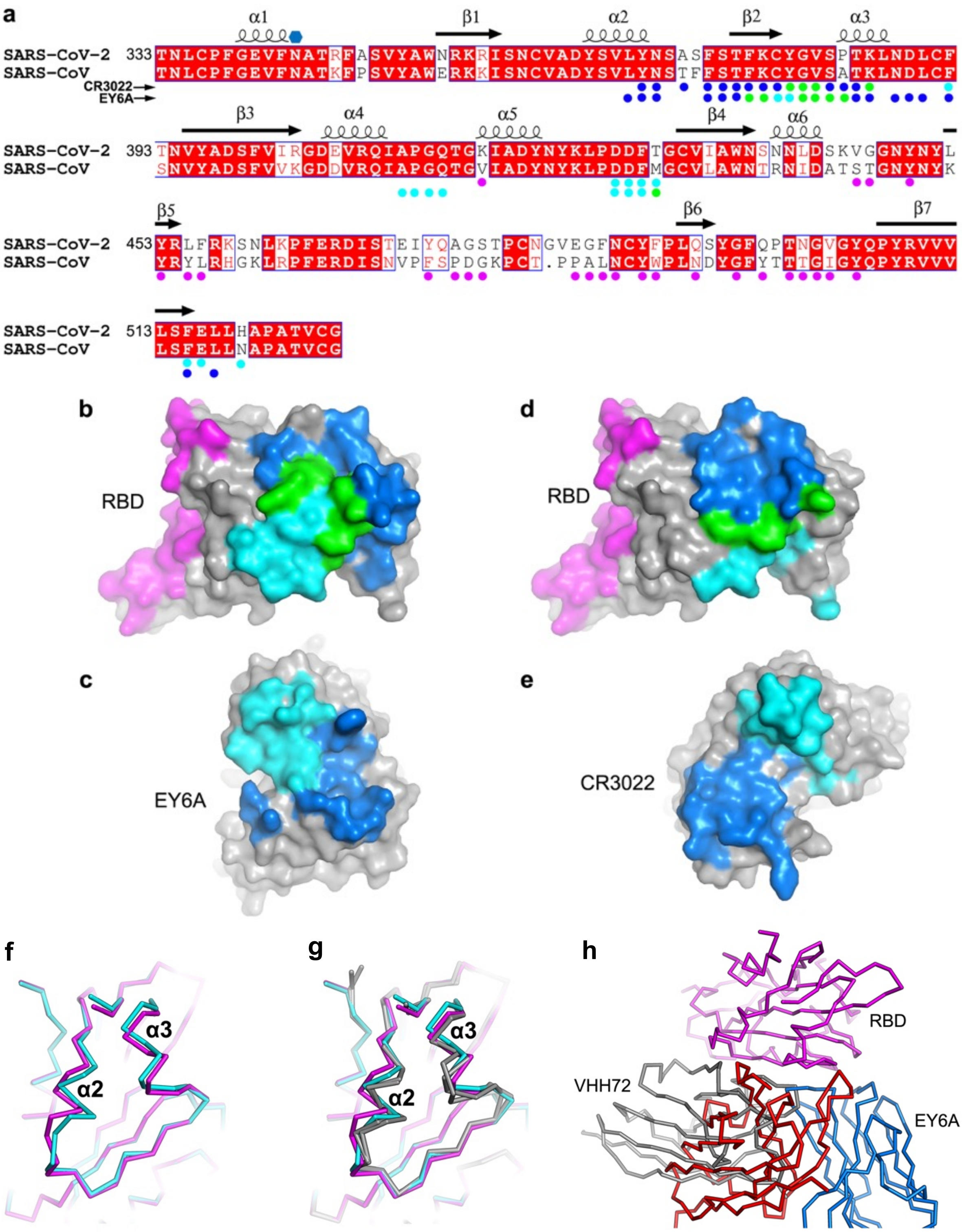


**Extended Data Fig. 6 | Comparison of EY6A, ACE2, CR3022 and epitopes on the RBDs of SARS-CoV-1 and SARS-CoV-2. a,** Sequence alignment of RBDs of SARS-CoV-2 and SARS-CoV-1. Residue numbers are those of SARS-CoV-2 RBD, conserved amino acids have a red background, secondary structures are labelled on the top of the sequence, and the glycosylation site is marked with a blue hexagon. Residues involved in receptor binding are marked with magenta disks. Residues shielded by Fab binding are marked with disks: blue indicate heavy chain, cyan light chain and green both chains. **b-e**, Open book views showing buried solvent accessible surface due to RBD-EY6A complex formation (**b, c**) and RBD-CR3022 complex formation (**c, d**). The colour scheme is as in (**a**). **f**, Superposition of EY6A and CR3022 bound RBDs showing the structural differences at the epitope region. EY6A bound RBD is shown in magenta and CR3022 bound RBD in cyan. **g**, Structural differences between EY6A and CR3022 bound RBDs and ACE2 bound RBDs (grey; PDB IDs, 6M0J and 6LZG). **h**, Comparison of binding modes between EY6A (red, heavy chain; blue, light chain) and VHH72 (grey) in the crystal structure of SARS-CoV-1 RBD/VHH72 complex (PDB ID, 6WAQ).

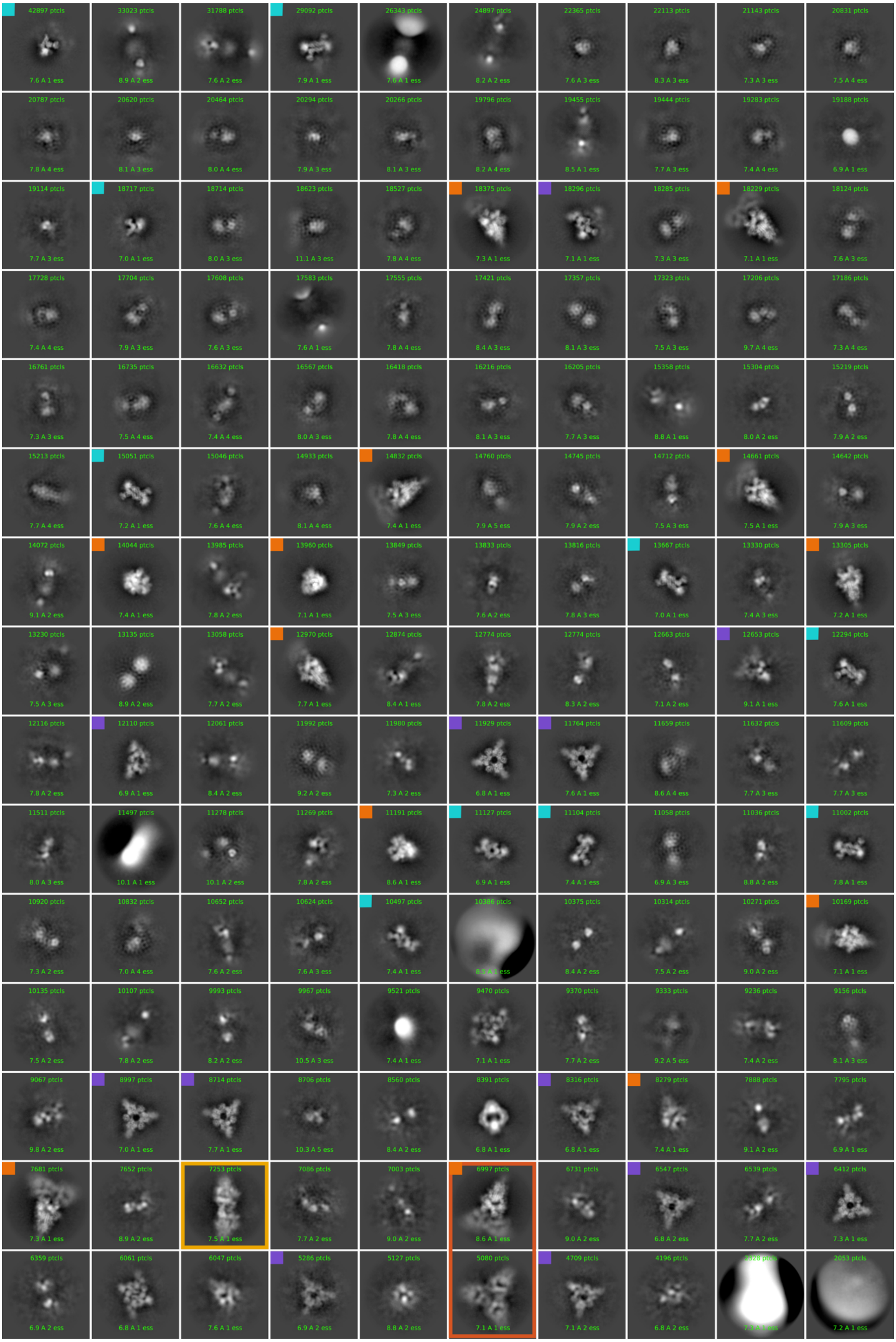
42897 ptcls
7.6 A 1 ess
33023 ptcls
8.9 A 2 ess
31788 ptcls
7.6 A 2 ess
29092 ptcls
7.9 A 1 ess
26343 ptcls
7.6 A 1 ess
24897 ptcls
8.2 A 2 ess
22365 ptcls
7.6 A 3 ess
22113 ptcls
8.3 A 3 ess
21143 ptcls
7.3 A 3 ess
20831 ptcls
7.5 A 4 ess
20787 ptcls
7.8 A 4 ess
20620 ptcls
8.1 A 3 ess
20464 ptcls
8.0 A 4 ess
20294 ptcls
7.9 A 3 ess
20266 ptcls
8.1 A 3 ess
19796 ptcls
8.2 A 4 ess
19455 ptcls
8.5 A 1 ess
19444 ptcls
7.7 A 3 ess
19283 ptcls
7.4 A 4 ess
19188 ptcls
6.9 A 1 ess
19114 ptcls
7.7 A 3 ess
18717 ptcls
7.0 A 1 ess
18714 ptcls
8.0 A 3 ess
18623 ptcls
11.1 A 3 ess
18527 ptcls
7.8 A 4 ess
18375 ptcls
7.3 A 1 ess
18296 ptcls
7.1 A 1 ess
18285 ptcls
7.3 A 3 ess
18229 ptcls
7.1 A 1 ess
18124 ptcls
7.6 A 3 ess
17728 ptcls
7.4 A 4 ess
17704 ptcls
7.9 A 3 ess
17608 ptcls
7.6 A 3 ess
17583 ptcls
7.6 A 1 ess
17555 ptcls
7.8 A 4 ess
17421 ptcls
8.4 A 3 ess
17357 ptcls
8.1 A 3 ess
17323 ptcls
7.5 A 3 ess
17206 ptcls
9.7 A 4 ess
17186 ptcls
7.3 A 4 ess
16761 ptcls
7.3 A 3 ess
16735 ptcls
7.5 A 4 ess
16632 ptcls
7.4 A 4 ess
16567 ptcls
8.0 A 3 ess
16418 ptcls
7.8 A 4 ess
16216 ptcls
8.1 A 3 ess
16205 ptcls
7.7 A 3 ess
15358 ptcls
8.8 A 1 ess
15304 ptcls
8.0 A 2 ess
15219 ptcls
7.9 A 2 ess
15213 ptcls
7.7 A 4 ess
15051 ptcls
7.2 A 1 ess
15046 ptcls
7.6 A 4 ess
14933 ptcls
8.1 A 4 ess
14832 ptcls
7.4 A 1 ess
14760 ptcls
7.9 A 5 ess
14745 ptcls
7.9 A 2 ess
14712 ptcls
7.5 A 3 ess
14661 ptcls
7.5 A 1 ess
14642 ptcls
7.9 A 3 ess
14072 ptcls
9.1 A 2 ess
14044 ptcls
7.4 A 1 ess
13985 ptcls
7.8 A 2 ess
13960 ptcls
7.1 A 1 ess
13849 ptcls
7.5 A 3 ess
13833 ptcls
7.6 A 2 ess
13816 ptcls
7.8 A 3 ess
13667 ptcls
7.0 A 1 ess
13330 ptcls
7.4 A 3 ess
13305 ptcls
7.2 A 1 ess
13230 ptcls
7.5 A 3 ess
13135 ptcls
8.9 A 2 ess
13058 ptcls
7.7 A 2 ess
12970 ptcls
7.7 A 1 ess
12874 ptcls
8.4 A 1 ess
12774 ptcls
7.8 A 2 ess
12774 ptcls
8.3 A 2 ess
12663 ptcls
7.1 A 2 ess
12653 ptcls
9.1 A 1 ess
12294 ptcls
7.6 A 1 ess
12116 ptcls
7.8 A 2 ess
12110 ptcls
6.9 A 1 ess
12061 ptcls
8.4 A 2 ess
11992 ptcls
9.2 A 2 ess
11980 ptcls
7.3 A 2 ess
11929 ptcls
6.8 A 1 ess
11764 ptcls
7.6 A 1 ess
11659 ptcls
8.6 A 4 ess
11632 ptcls
7.7 A 3 ess
11609 ptcls
7.7 A 3 ess
11511 ptcls
8.0 A 3 ess
11497 ptcls
10.1 A 1 ess
11278 ptcls
10.1 A 2 ess
11269 ptcls
7.8 A 2 ess
11191 ptcls
8.6 A 1 ess
11127 ptcls
6.9 A 1 ess
11104 ptcls
7.4 A 1 ess
11058 ptcls
6.9 A 3 ess
11036 ptcls
8.8 A 2 ess
11002 ptcls
7.8 A 1 ess
10920 ptcls
7.3 A 2 ess
10832 ptcls
7.0 A 4 ess
10652 ptcls
7.6 A 2 ess
10624 ptcls
7.6 A 3 ess
10497 ptcls
7.4 A 1 ess
10386 ptcls
10375 ptcls
8.4 A 2 ess
10314 ptcls
7.5 A 2 ess
10271 ptcls
9.0 A 2 ess
10169 ptcls
7.1 A 1 ess
10135 ptcls
7.5 A 2 ess
10107 ptcls
7.8 A 2 ess
9993 ptcls
8.2 A 2 ess
9967 ptcls
10.5 A 3 ess
9521 ptcls
7.4 A 1 ess
9470 ptcls
7.1 A 1 ess
9370 ptcls
7.7 A 2 ess
9333 ptcls
9.2 A 5 ess
9236 ptcls
7.4 A 2 ess
9156 ptcls
8.1 A 3 ess
9067 ptcls
9.8 A 2 ess
8997 ptcls
7.0 A 1 ess
8714 ptcls
7.7 A 1 ess
8706 ptcls
10.3 A 5 ess
8560 ptcls
8.4 A 2 ess
8391 ptcls
6.8 A 1 ess
8316 ptcls
6.8 A 1 ess
8279 ptcls
7.4 A 1 ess
7888 ptcls
9.1 A 2 ess
7795 ptcls
6.9 A 1 ess
7681 ptcls
7.3 A 1 ess
7652 ptcls
8.9 A 2 ess
7253 ptcls
7.5 A 1 ess
7086 ptcls
7.7 A 2 ess
7003 ptcls
9.0 A 2 ess
6997 ptcls
8.6 A 1 ess
6731 ptcls
9.0 A 2 ess
6547 ptcls
6.8 A 2 ess
6539 ptcls
7.7 A 2 ess
6412 ptcls
7.3 A 1 ess
6359 ptcls
6.9 A 2 ess
6061 ptcls
6.8 A 1 ess
6047 ptcls
7.6 A 1 ess
5286 ptcls
6.9 A 2 ess
5127 ptcls
8.8 A 2 ess
5080 ptcls
7.1 A 1 ess
4709 ptcls
7.1 A 2 ess
4196 ptcls
6.8 A 2 ess
2053 ptcls
7.2 A 1 ess

**Extended Data Fig. 7 | 2D class averages for 5h incubation.** Unbiased 2D classes from blob-picked particles in CryoSPARC. The three major particle forms are indicated by coloured boxes located at the top left of each box; Population A (purple, 'trimeric association'): 153,502 particles; Population B (cyan, 'dimeric association'): 103,623 particles, Population C (orange, pre-fusion Spike): 175,448 particles. See Methods for details. In some classes, boxed in pale and dark orange, Spike oligomers were seen, where the Fab linked two Spike heads together.

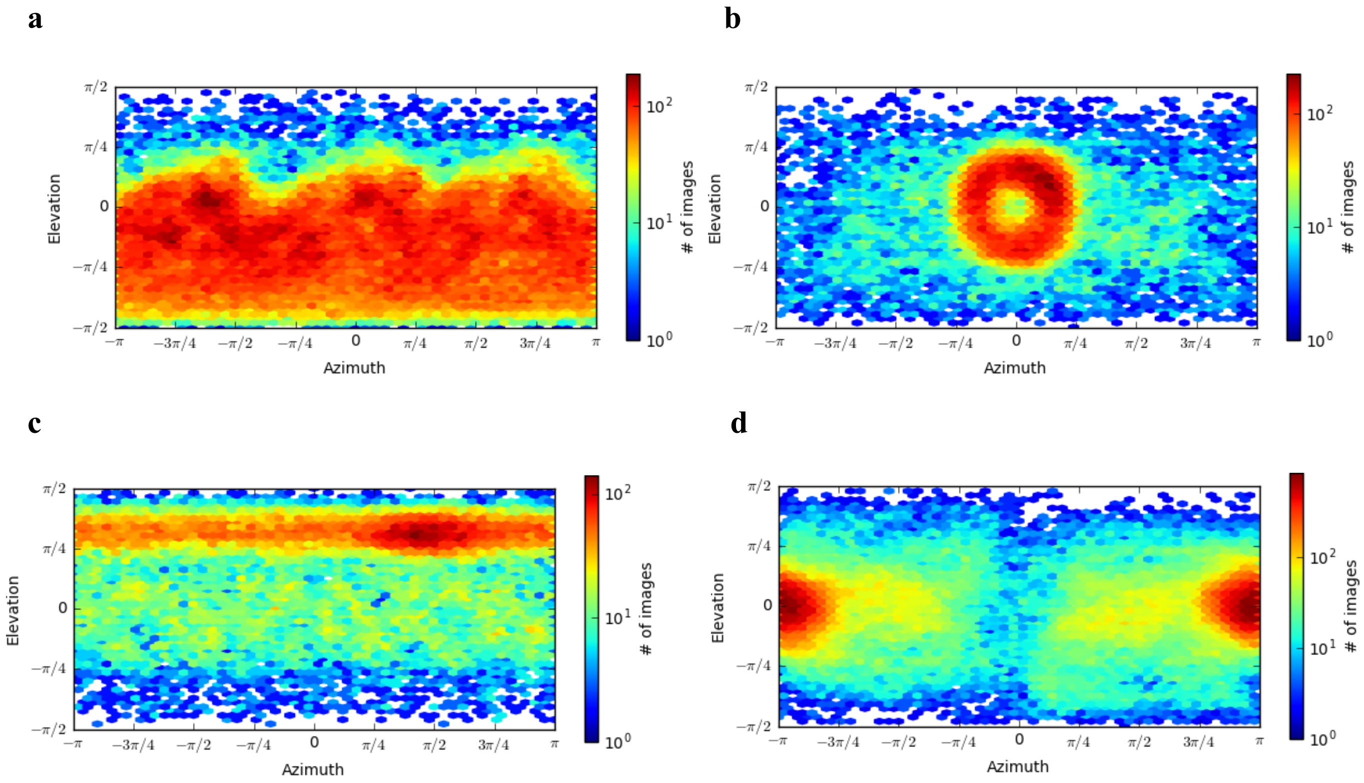


**Extended Data Fig. 8 | Cryo-EM Structure view direction distribution plots. a,** EY6A-bound intact Spike; **b,** EY6A-RBD ring with C1 and **c** with C3 (ii) symmetry imposed**. d,** EY6A-Spike dimer. Plots were generated within CryoSPARC. See Methods for details

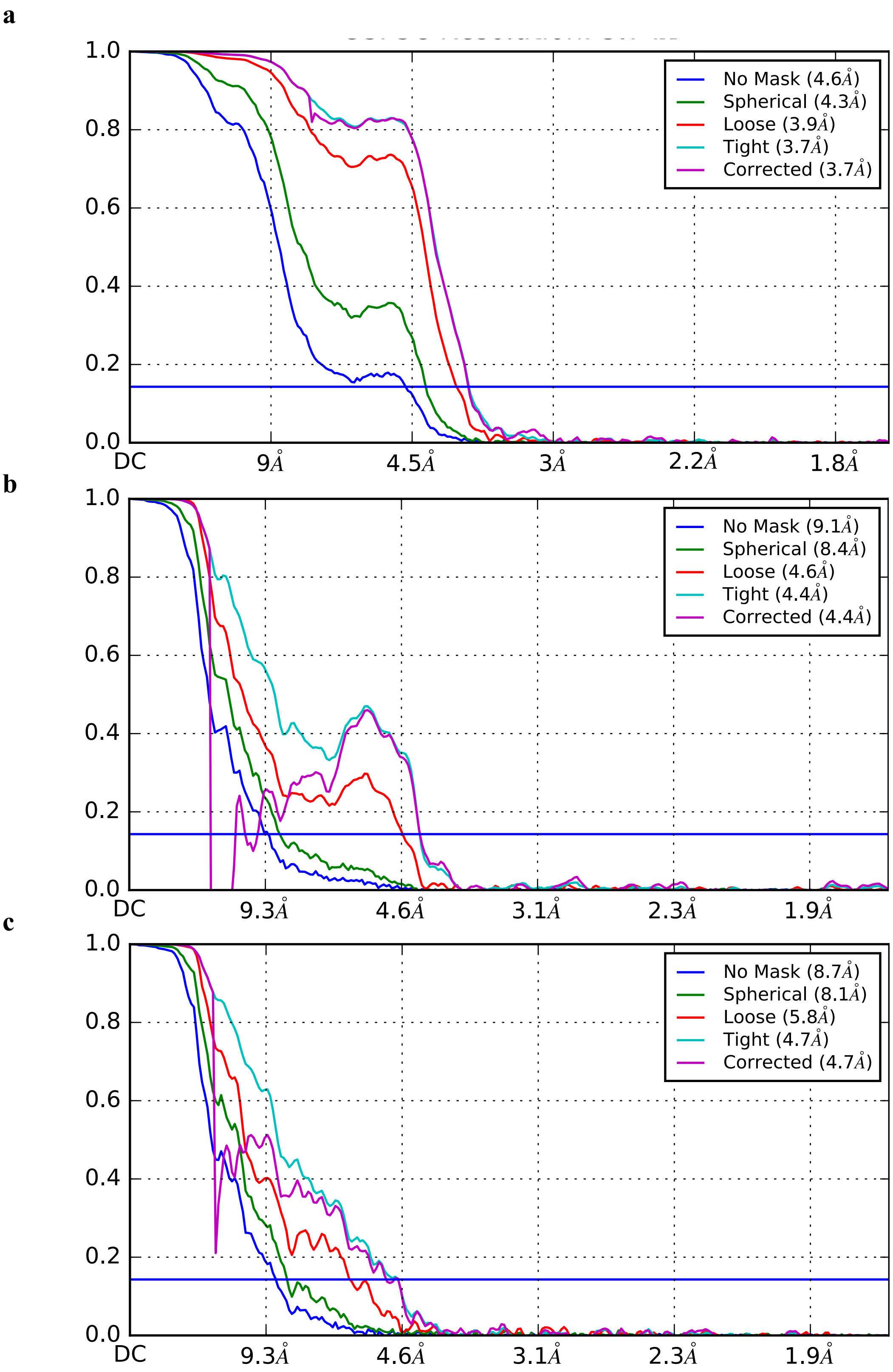


**Extended Data Fig. 9 | FSC curves for cryo-EM reconstructions. a,** EY6A-bound intact Spike; **b,** ‘dimeric fragment’; **c**, ‘trimeric fragment’

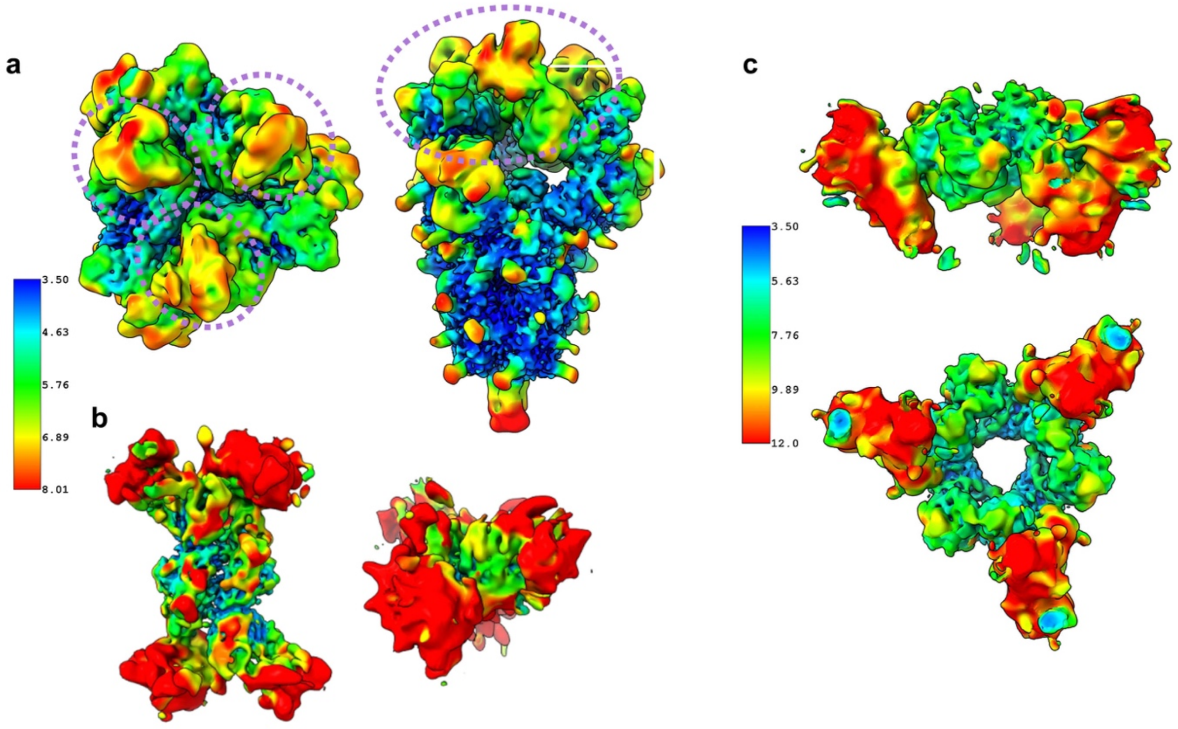


**Extended Data Fig. 10 | Resolution maps for the three cryo-EM reconstructions.** **a** pre-fusion trimer (the purple dotted lines mark the RBD positions). **b** ‘dimeric assembly’. **c** ‘trimeric assembly’

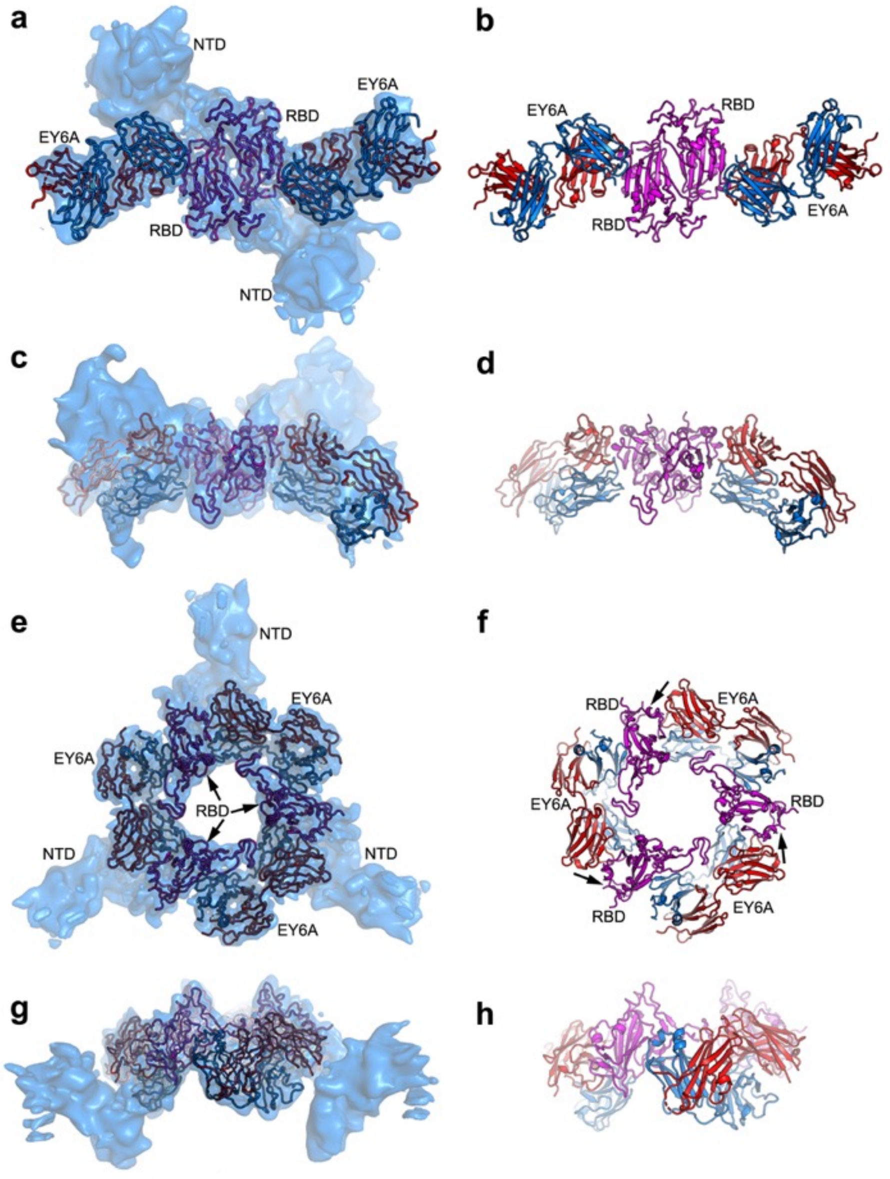


**Extended Data Fig. 11 | Analysis of cryo-EM data for 5 h incubation.** Final reconstructions derived using an *ab initio* CryoSPARC reference volume from the 5 h incubation dataset to show two observed degraded states of Spike-EY6A Fab complex. **a, b,** Top view **and c, d,** side view of the 'dimeric' assembly. **e, f,** Top view and **g, h** side view of the 'trimeric' assembly. EY6A is drawn as a cartoon in blue (light chain) and red (heavy chain), and the RBD in magenta.

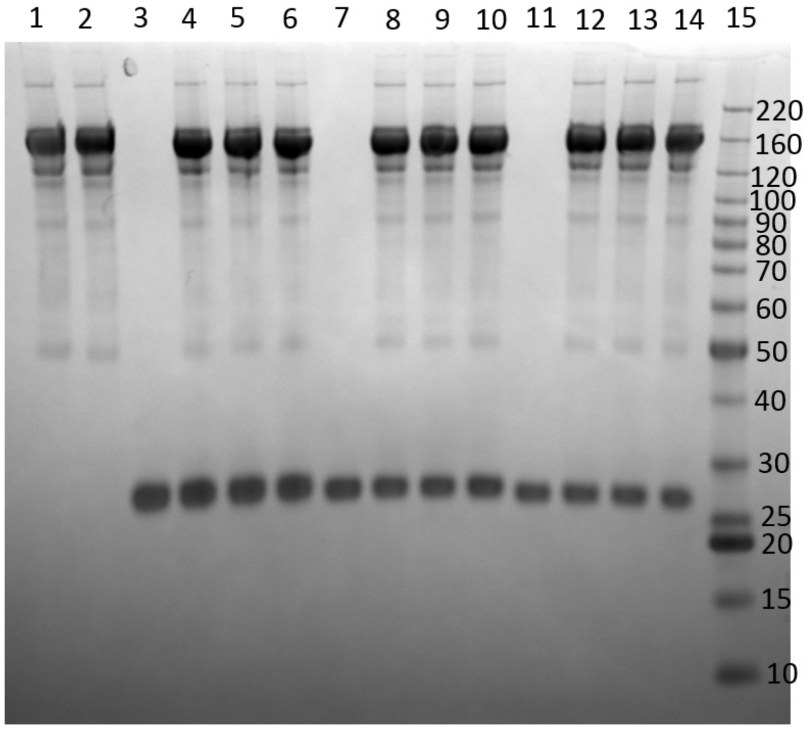


**Extended Data Fig. 12 | SDS-PAGE analysis of material following 5 h incubation, as for cryo-EM.** The gel is a 4-12% gradient SDS-PAGE gel (under reducing conditions). Lanes 1 & 2 are Spike alone. Lane 3 is CR3022 Fab alone, lanes 4-6 are Spike incubated with CR3022 Fab. Lane 7 is EY6A Fab alone, lanes 8-10 are Spike incubated with EY6A Fab. Lanes 11-14 are a non-RBD bindig Fab alone and incubated with Spike. Lane 15 is molecular weight markers (kDa).

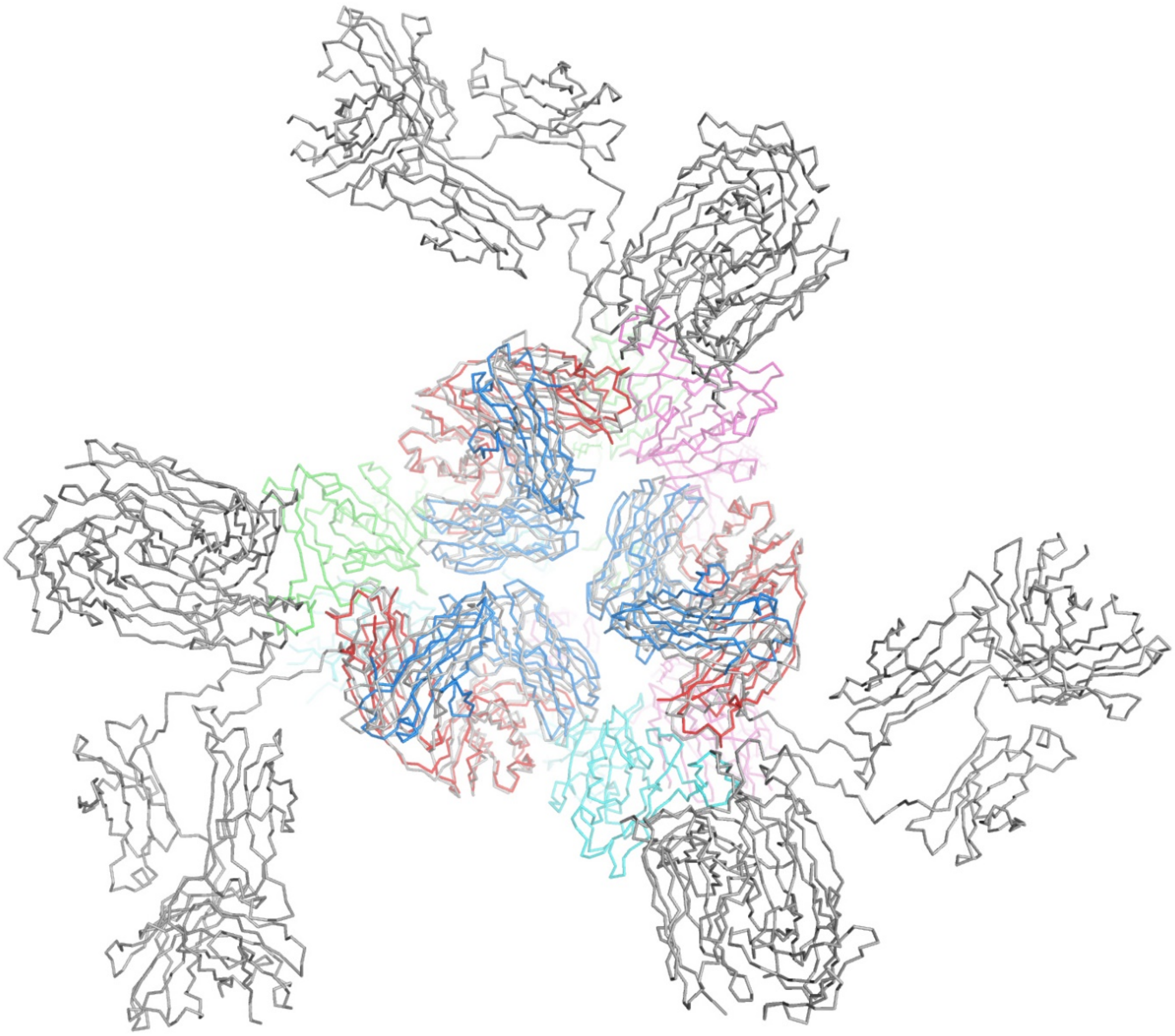

**Extended Data Fig. 13 | Intact EY6A antibodies can pack similarly to EY6A Fabs at the pre-fusion Spike head.** Representative antibody structures (PDB ID, 1IGT) have been superposed on the EY6A Fab without significant clashes. The Spike chains are coloured green, cyan and magenta with EY6A heavy and light chains shown in red and blue respectively and the remainder of the antibodies in grey.

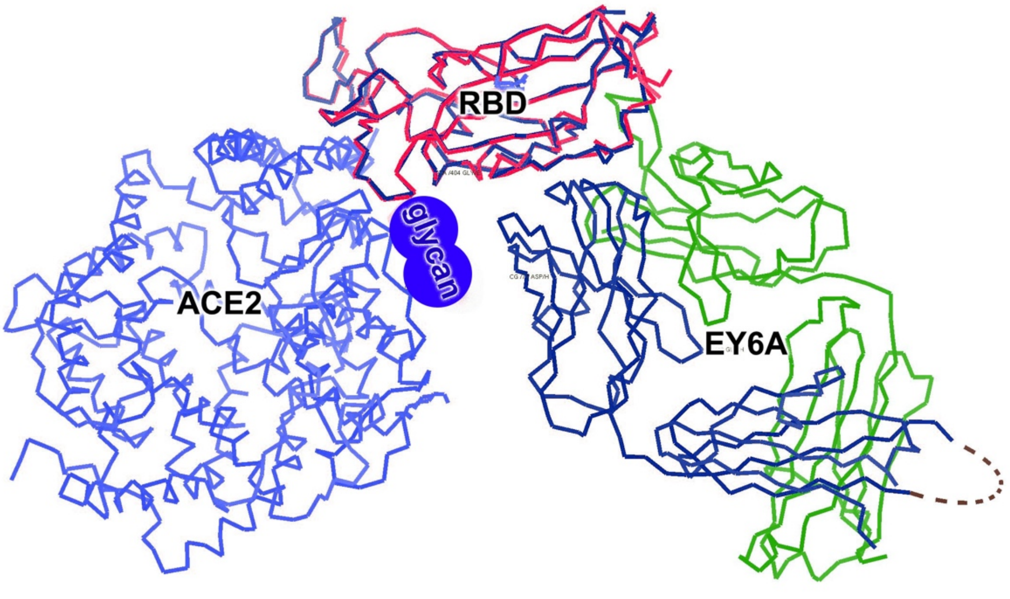


**Extended Data Fig. 14 | Glycans may modulate interactions between ACE2 and EY6A.**

**Extended Data Table 1 SPR Average kinetic results**

| Ligand | RBD - Fc | EY6A IgG |
|---|---|---|
| Analyte | EY6A Fab | RBD |
| Ka ($M^{-1}s^{-1}$) | 5.3E+05 | 6.0E+05 |
| Kd ($s^{-1}$) | 1.0E-03 | 4.6E-03 |
| $K_D$ (nM) | 2 | 8 |

**Extended Data Table 2 Plaque Reduction Neutralization Test results (PHE, Porton Down)**

| ID | Description | ND50 |
|---|---|---|
| Positive control | MERS Convalescent serum | 1:784 |
| EY6A | Mab | 1:251 |

**Extended Data Table 3 X-ray data collection and refinement statistics**

| **Data collection** | | |
|---|---|---|
| Data set | RBD-EY6A | RBD-EY6A-Nb |
| Space group | *$P3_121$* | *R3* |
| Cell dimensions | | |
| *a, b, c* (Å) | *166.6, 166.6, 270.8* | *178.1, 178.1, 87.8* |
| *α, β, γ* (°) | *90, 90, 120* | *90, 90, 120* |
| Resolution (Å) | 144–3.80 (3.87–3.80) | 89–2.64 (2.69–2.64) |
| Unique reflections | 43446 (2141) | 30147 (1419) |
| $R_{\mathrm{merge}}$ | 0.227 (---) | 0.209 (---) |
| $R_{pim}$ | 0.052 (0.636) | 0.071 (1.369) |
| $CC_{1/2}$ | 0.998 (0.783) | 0.993 (0.298) |
| $\langle I \rangle / \langle \sigma I \rangle$ | 7.3 (0.4) | 5.0 (0.20) |
| Completeness (%) | 100 (100) | 99.2 (93.0) |
| Redundancy | 19.8 (19.8) | 9.4 (5.3) |
| **Refinement** | | |
| Resolution (Å) | 28.52–3.80 | 35.3–2.65 |
| No. reflections (work/test) | 40960/2156 | 25517/1267 |
| $R_{\mathrm{work}}$ / $R_{\mathrm{free}}$ | 0.219/0.259 | 0.216/0.262 |
| No. atoms | 14520 | 5837 |
| Average *B*-factors ( $Å^2$) | 184 | 82 |
| R.m.s. deviations | | |
| Bond lengths (Å) | 0.003 | 0.002 |
| Bond angles (°) | 0.7 | 0.4 |

Numbers in brackets refer to the highest resolution shell of data

**Extended Data Table 4 Cryo-EM data collection and refinement parameters**

| | (A) SARS-CoV-2 Spike | (B) S1-EY6A trimer | (C) RBD-EY6A dimer |
|---|---|---|---|
| **Data collection and reconstruction** | | | |
| Voltage (kV) | 300 | | |
| Frames | 40 (50) | | |
| Dose rate ($e^-$/ Å $^2$/ s) | 20.0 (20.0) | | |
| Total dose ($e^-$/ Å $^2$) | 42.2 (52.5) | | |
| Pixel size (Å) | 0.415 super-resolution | | |
| Defocus (μm) | 0.8-2.6 | | |
| Symmetry | C1 | C3 | C1 |
| Movies | 6420 (6129) | | |
| Particles | 144,680 | 41,372 | 119,343 |
| Refined Map resolution FSC = 0.5 (Å) | 3.7 | 4.7 [5.9] | 4.4 |
| Map sharpening B-factor (Å$^2$) | -91 | -206 [-298] | -154 |
| **Model refinement** | | | |
| Model-to-map fit, CC_mask | 0.72 | 0.46 | 0.33 |
| R.m.s.d., bonds (Å) | 0.005 | 0.005 | 0.002 |
| R.m.s.d., angles (°) | 0.6 | 0.6 | 0.5 |
| All-atom Clash score | 5.6 | 9.3 | 6.5 |
| Rotamer outliers (%) | 1.1 | 1.5 | 1.5 |
| Ramachandran plot | | | |
| Favoured (%) | 96.6 | 92.4 | 92.6 |
| Allowed (%) | 3.4 | 7.6 | 7.4 |
| Outliers (%) | 0 | 0 | 0 |

Numbers in brackets refer to the 30º tilted dataset that was merged with the 0º data. Square brackets provide values for C1 symmetry.
